# A taxonomic review and revisions of Microstomidae (Platyhelminthes: Macrostomorpha)

**DOI:** 10.1101/381459

**Authors:** Sarah Atherton, Ulf Jondelius

## Abstract

Microstomidae (Platyhelminthes: Macrostomorpha) diversity has been almost entirely ignored within recent years, likely due to inconsistent and often old taxonomic literature and a general rarity of sexually mature collected specimens. Herein, we reconstruct the phylogenetic relationships of the group using both previously published and new 18S and CO1 gene sequences. We present some taxonomic revisions of Microstomidae and further describe 8 new species of *Microstomum* based on both molecular and morphological evidence. Finally, we briefly review the morphological taxonomy of each species and provide a key to aid in future research and identification that is not dependent on reproductive morphology. Our goal is to clarify the taxonomy and facilitate future research into an otherwise very understudied group of tiny (but important) flatworms.

## Introduction

Macrostomorpha DOE, 1986 is a group of free-living Platyhelminthes found worldwide in aquatic and semi-aquatic habitats (Rieger 2001). They are very small in size, typically only 1-2 mm, but are nevertheless important and often abundant members of their communities, feeding on diatoms and other microorganisms or even other small invertebrates (Ax 1995; Carrasco 2012; Atherton & Jondelius 2018b). While the deep splits within Macrostomorpha remain ambiguous, current phylogenetic hypotheses (Littlewood et al. 1999; Litvaitis & Rohde, 1999; Janssen et al. 2015), place Microstomidae as sister to Dolichomacrostomidae within the group Dolichomicrostomida.

Microstomidae LUTHER 1907 are characterised by ciliated sensory pits at the frontal end, an intestine extending anteriorly past the mouth, and the ability to reproduce asexually through fissioning. Currently, Microstomidae is unevenly divided between three genera: *Myozonella* BEKLEMISHEV, 1955, with only a single species *Myozonella microstomoides* BEKLEMISHEV, 1955 collected from a village in western Russia and characterized by a muscular ring surrounding the anterior intestine; *Alaurina* BUSCH 1851, with three currently accepted species all with an elongated and unciliated (apart from sensory cilia) anterior proboscis; and the highly diverse and speciose genus *Microstomum* ØRSTED 1843.

Species of Microstomidae have been considered difficult to distinguish through traditional morphological methods. Their small size, relatively few morphological characters, and high amounts of intraspecific variation (Luther 1960; Bauchhens 1971; Heitkamp 1982; Atherton & Jondelius 2018b) all present taxonomic challenges, especially as sexually mature specimens are generally rare. Furthermore, the taxonomic literature may sometimes lack detail and is often old and/or difficult to access, such that even the otherwise-reliable WORMS database (http://www.marinespecies.org/aphia.php?p=taxdetails&id=142210) includes numerous mistakes (e.g. recognition of *M. hanatum*, still listing *M. giganteum* as synonymous to *M. lineare*). Taken together, it is unsurprising that while there has been some recent studies of species diversity within Macrostomorpha in general (e.g., Schockaert 2014; Sun et al. 2014; Janssen et al. 2015; Fang et al. 2016; Reyes & Brusa 2017; Lin et al. 2017a, b;), taxonomic research within Microstomidae from even the last 50 years has been restricted to a very few articles (Faubel 1974, 1984; Rogozin 2015; Atherton & Jondelius 2018 a,b).

Species of Microstomidae primarily reproduce through asexual fission, but their life cycle may also include short periods of sexual reproduction occurring primarily in the autumn (Bauchhenss 1971; Heitkamp 1982). During this time, individual zooids will develop male and female sexual structures, including for the male complex: single or bilateral testes, vasa deferentia, and a male copulatory apparatus with vesicula seminalis, stylet and antrum masculinum; and for the female complex: a single ovary and a female antrum. Species of Microstomidae may be sexually mature for as little as two weeks a year or less (Bauchhenss 1971; Faubel 1974) and thus full or partial sexual anatomy is currently only known for roughly half of the nominal species. Since species identification of Macrostomorpha has often been based heavily or, in some cases, entirely on the morphology of the reproductive organs (e.g. Nasonov 1935; Ax & Armonies 1987; Faubel & Cameron 2001), some authors have argued that any species with unknown sexual anatomy should be instantly considered species dubiae (Faubel 1984).

For Microstomidae with limited morphological data available, analysis of nucleotide sequence data will be an indispensable tool when solving taxonomic problems. DNA taxonomy is not without its own pitfalls (e.g. sensitivity to sampling, overlap between intra and interspecific variation; DeSalle et al. 2005; Meier et al. 2006; Sauer & Hausdorf 2012), but has nevertheless been demonstrated as a reliable and reproducible method to identify lineages without depending solely on morphological characters that may be sensitive to environmental conditions or access to sexually mature specimens (Pérez-Ponce de León & Poulin 2016; Fonseca et al. 2017). A number of different approaches to sequence-based species discovery, or DNA-taxonomy, have been developed, including haplotype networks based on statistical parsimony (Templeton *et al.* 1992), Bayesian species delineation (Yang & Rannala 2010; Zhang *et al.* 2013), and maximum likelihood approaches with the General Mixed Yule-Coalescent model (Pons *et al.* 2006).

DNA taxonomy may be more effective than traditional morphological methods in discerning species, as re-examination of morphological taxa often reveals hidden biodiversity in morphologically inseparable, i.e. cryptic species (Fiser et al. 2018). Successful DNA-taxonomy requires use of a molecular marker with an appropriate level of variation, or, preferably, more than one such marker (Fontaneto et al. 2015). Recent molecular investigations into the widespread taxon *Microstomum lineare* revealed an assemblage of four genetically divergent but morphologically similar species, some of which were found in the same sample (Atherton & Jondelius 2018b). The presence of multiple distinct but cryptic lineages in single locations potentially could have much larger implications for Microstomidae research, especially since studies in the past have often been performed on specimens collected from environmental samples (e.g. Bauchhenss 1971; Heitkamp 1982; Reuter & Palmberg 1989; Gutafsson et al. 1995).

In this paper, we reconstruct the phylogenetic relationships of *Microstomum* using both previously published and new 18S and CO1 nucleotide sequences. Our dataset comprises sequences from 336 specimens representing 15 previously described nominal species and eight new species. The new taxa are diagnosed on the basis of DNA-taxonomy as well as morphological characters that distinguish them from all other known *Microstomum* species. We also review the morphological taxonomy of the species within Microstomidae and provide a key to aid in future research and identification. Our goals are to simplify the taxonomy, demonstrate that asexual *Microstomum* specimens can often be identified, and to facilitate future research into an otherwise very understudied group of microscopic flatworms that are highly abundant in both limnic and marine habitats.

## Methods

Sediments and aquatic vegetation were collected from fresh and marine waters. Collection details for each specimen, including sampling date and GPS coordinates, are listed in Table S1. Samples were transported back to laboratories and processed according to type. Animals were extracted from marine sediments following a standard anesthetization-decantation technique (Martens 1984) within 48 hours of collection. Freshwater sediments were first allowed to stagnate for 24-72 hours before the water was either directly removed into a glass petri dish or was first filtered through a 63 μm sieve and backwashed to the petri dish. Vegetation was simply washed through a fine mesh sieve.

Following extraction, animals were manually isolated under a Nikon SMZ 1500 stereomicroscope, transferred to a glass slide and identified with either a Nikon Eclipse 80i compound microscope equipped with DIC (differential interference contrast) or a Leitz LaborLux S compound microscope. Light micrographs and digital videos were captured with a Cannon EOS 5D Mark III digital camera, and measurements were taken with an ocular micrometer. Following documentation, individual specimens were fixed in 95% ethanol and transported to Naturhistoriska riksmuseet in Stockholm for DNA extraction and analysis.

DNA was extracted from whole animals using the DNeasy Blood & Tissue kit (Qiagen, Valencia, CA) following the manufacturer’s instructions. PCR amplification was performed using 0.2 ml PuReTaq Ready-To-Go PCR Beads (GE Healthcare) with 5 pmol each forward and reverse primers and 2 μl DNA. Table S2 lists primer pairs and protocols used for amplification and sequencing. Products were viewed on 1-2% agarose gels, purified using ExoSAP-IT enzymes (Exonuclease and Shrimp Alkaline Phosphatase; GE Healthcare) and sent for commercial sequencing to Macrogen Europe (Netherlands) for commercial sequencing.

Amplification was performed on the complete nuclear 18S gene as well as a ~650 bp segment of the mitochondrial cytochrome oxidase c subunit 1 (CO1) loci pertaining to the “Folmer region” (Folmer et al. 1994). The 18S ribosomal gene is currently the most commonly used molecular marker for flatworms on GenBank (8895 entries as of 5 July 2018), while CO1 gene sequences are known to be highly variable in Platyhelminthes (e.g. Larsson et al. 2008; Telford et al. 2000; Atherton & Jondelius 2018b) and other meiofauna (e.g. Verheye et al. 2016; Álvarez-Campos et al. 2017; Kieneke & Nikoukar 2017; Michaloudi et al. 2017; Sahraean et al. 2017; Zhao et al. 2018) and are thus applicable in identifying species and assessing intraspecific diversity.

Two catenulid species, *Stenostomum leucops* and *Parcatenula* sp., and one polyclad species, *Hoploplana californica*, were selected as outgroup taxa. Catenulida is considered the sister group of all Rhabditophora, and Polycladida is the closest relative of the Macrostomorpha (Egger et al. 2015; Laumer et al. 2015). Outgroup sequences as well as available species of Macrostomorpha were downloaded from GenBank and combined with new sequences for analysis. Table S1 list accession numbers for all specimens used in subsequent analyses

Digital morphological vouchers (pictures and video) as well as collection data of four previously undescribed *Microstomum* species were downloaded from the Dryad Digital Repository (https://datadryad.org/resource/doi:10.5061/dryad.b5908) and the Macrostomorpha Taxonomy and Phylogeny website (http://www.macrostomorpha.info/). Further information on the collection and processing of these species can be found in Janssen et al. (2015).

Sequence assembly and alignment was performed in Mega v 7.0.21 (Kumar et al. 2016), and trace files were manually edited. Alignments for each marker were generated using MUSCLE (Edgar 2004) with the default settings. CO1 sequences were translated to amino acids using the flatworm mitochondrial genetic code (Telford et al. 2000), manually checked for stop codons and reading frame shifts, aligned and then reverted back to the original nucleotides for analysis. Maximum likelihood (ML) analysis was performed on each marker individually along with concatenated datasets in IQTree v1.6.4. The best fitting model was determined using the ModelFinder algorithm (Kalyaanamoorthy et al. 2017) implemented in IQTree. We performed 1000 ultrafast bootstrap replicates (Nguyen et al. 2015) in IQTree, applying edge-proportional partitions with best-fit models independently selected for each gene in the concatenated datasets.

Species of Microstomidae were inferred using the Bayesian implementation of the Poisson Tree Processor (bPTP) in the web-based interface (http://species.hits.org/) with the default parameters (Zhang et al. 2013). Input trees were generated from ML analyses of concatenated datasets that included Microstomidae sequences and two sequences each of *Myozonaria bistylifera*, *Myozonaria fissipara* and otherwise unidentified Myozonariinae downloaded from GenBank as outgroups (Table S1). All bPTP tests were re-run up to three times with the input tree pruned so that each prospective species included a maximum of five randomly selected sequences in order to ensure that results were not influenced by an imbalance in the number of specimens in any given clade. Following recommended guidelines, outgroups were not included in any bPTP analysis.

## Results

The full-concatenated alignment had a total of 336 specimens and total length of 2348 bp. The K3Pu+F+G4 substitution model was selected as the best fit for the CO1 gene sets and the TIM2e+I substitution model was selected for the 18S gene sets. Tree topologies were consistent across all individual gene and concatenated analyses.

Fig. 1 summarizes the results of the entire concatenated phylogeny (see also Fig. S1). Results were generally consistent with Janssen et al. (2015), with maximum (1.00) support for the two major subgroupings Macrostomidae and Dolichomicrostomida. Results found further high (>0.90) support for: a monophyletic Macrostomidae consisting of *Macrostomum* and *Psammomacrostomum*; Myozonariinae as sister to Dolichomacrostominae, including the genera *Paramalostomum*, *Austromacrostomum*, *Cylidromacrostomum* and *Dolichomacrostomum*; and a monophyletic Microstomidae, which in this analysis included *Alaurina composita* nested within *Microstomum*.

**Fig. 1:**
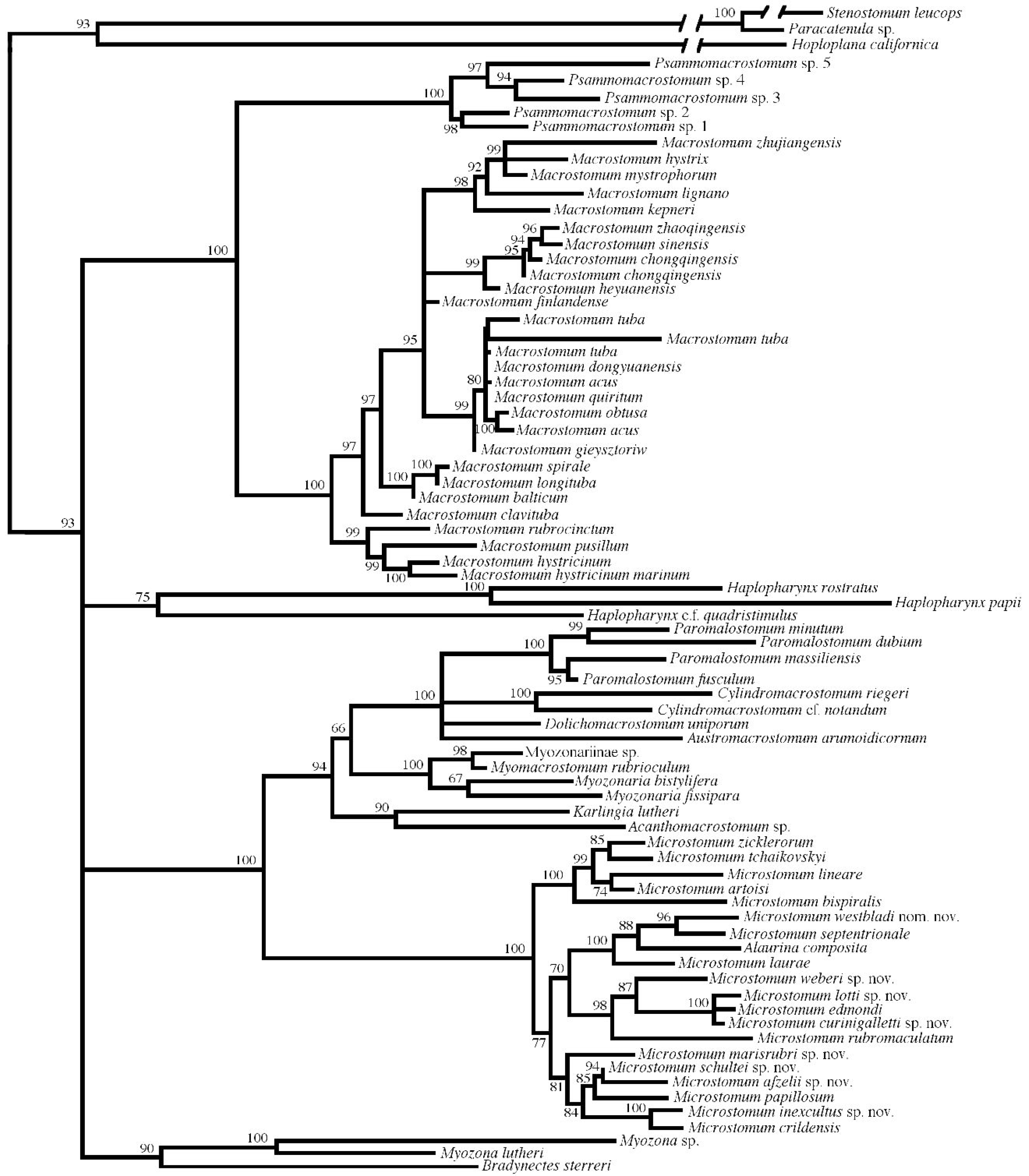
Concatenated 18S and CO1 gene tree summary. Percent bootstrap values are given at each node.

bPTP results were consistent across analyses, and a total of twenty species were identified within the *Microstomum* clade (Fig. 2). Twelve of these were previously described: *Alaurina composita* METSCHNIKOV 1865, *Microstomum artoisi* ATHERTON & JONDELIUS 2018, *M. bispiralis* STIREWALT 1937, *M. crildensis* FAUBEL 1984, *M. edmondi* ATHERTON & JONDELIUS 2018, *M. laurae* ATHERTON & JONDELIUS 2018, *M. lineare* (MÜLLER 1773), *M. papillosum* (GRAFF 1882), *M. rubromaculatum* (GRAFF 1882), *M. septentrionale* (SABUSSOW 1900), *M. tchaikovskyi* ATHERTON & JONDELIUS 2018, and *M. zicklerorum* ATHERTON & JONDELIUS 2018. The other eight species are new and are described below.

**Fig. 2:**
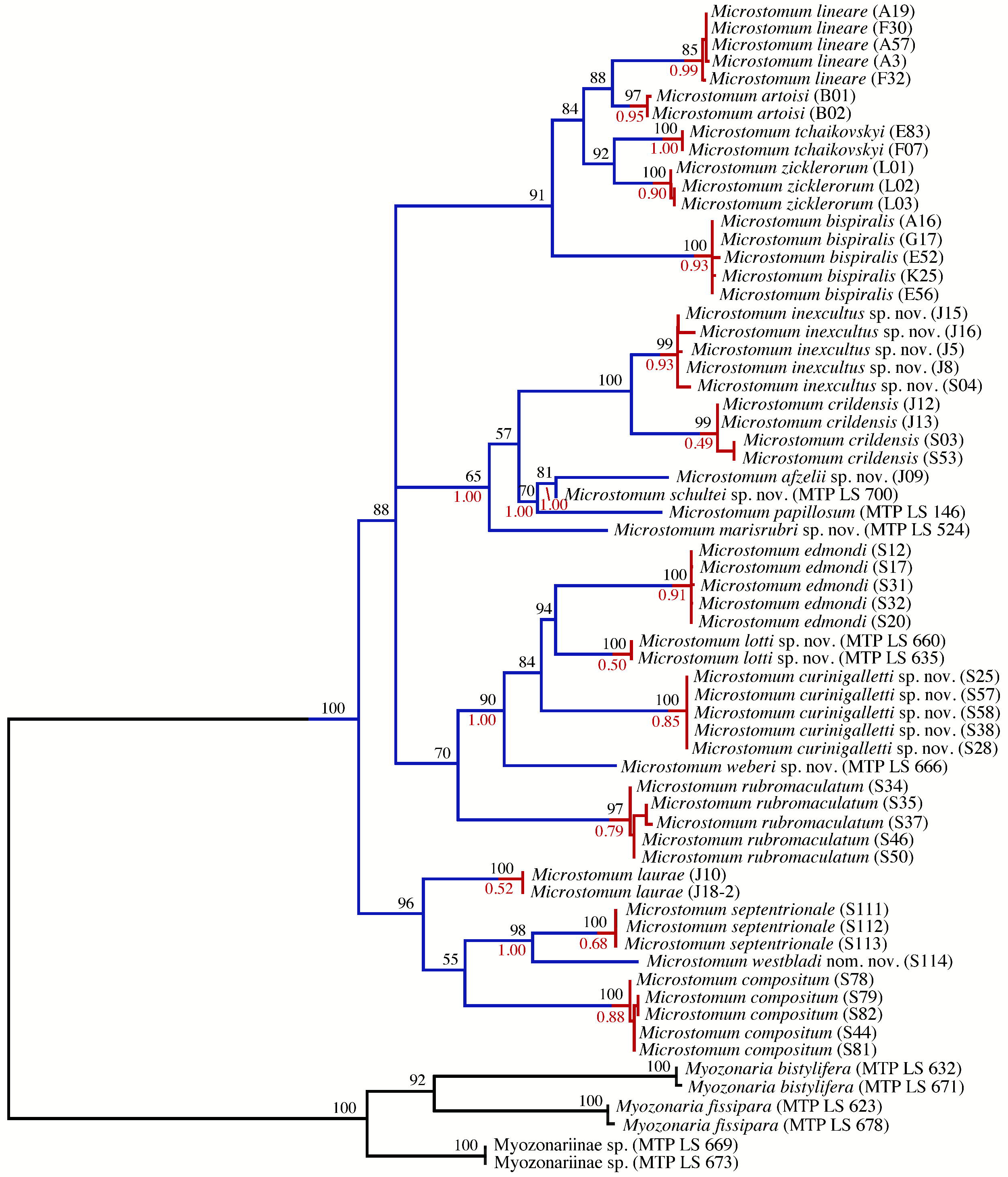
bPTP analysis of 18S and CO1 gene tree. A maximum of five specimens were chosen for each species. Specimens are listed with the isolate label in parentheses. At each node, percent bootstrap values (black above) and bPTP (red below) values are given. Transitions from blue to red lines highlight species according to the results of the bPTP analysis.

We obtained high support for four main clades within *Microstomum* (Figs. 1, 2), composed of: 1. all five freshwater species included in the analysis (*M. artoisi*, *M. bispiralis*, *M. lineare*, *M. tchaikovskyi*, *M zicklerorum*); 2. six species (*M. afzelli* sp. nov., *M. crildensis*, *M. inexcultus*, *M. marisrubri* sp. nov., *M. papillosum, M. schultei* sp. nov.); 3. five species (*M. curinigalletti* sp. nov., *M. edmondi*, *M. lotti* sp. nov., *M. rubromaculatum*, *M. weberi* sp. nov); and 4. *Alaurina composita* grouping with *M. laurae*, *M. septentrionale* and *M. westbladi* nom.

Unfortunately, nucleotide sequences from two species of *Alaurina* and one species of *Myozonella* were unavailable due to lack of specimens; however, results from our phylogenetic analyses found all specimens of *Alaurina composita* nested within the *Microstomum* clade (Figs. 1, 2), and thus we herein propose to synonymise *Alaurina* with *Microstomum* and transfer its three species to *Microstomum*.

## Nomenclature Acts

*Synonymization of* Alaurina *and amended diagnosis of* Microstomum *SCHMIDT 1848*

## Gen. Microstomum Schmidt 1848

Microstomidae with ciliary pits at frontal end and preoral intestine. Two excretion openings in the anterior body. Rhabdites and ingested nematocysts present in some species. Asexual reproduction through fissioning as well as sexual reproduction. Sexual anatomy including one or two testes beside or behind the intestine; one or two ovaries consisting of one or more lobes and containing a central oocyte and a peripheral layer of nutrient cells. Body size of solitary individual 0.6-4mm, with chains comprising up to 18 zooids and 15 mm.

Type species: *Microstomum lineare* (Müller, 1773).

43 species: 33 marine, 9 limnic, and 1 inhabiting both brackish (up to 7‰) and freshwaters.

*Synonymization of* Microstomum hamatum *WESTBLAD 1953 and* Microstomum lucidum *(FUHRMANN, 1896)*

*M. lucidum* and *M. hamatum* are morphological identical accepting the ambiguously-defined “dark greyish ± pigment spots” in *M. hamatum* (Westblad 1953). Both species have bodies colored yellow due to the intestine, with identical body shapes and sizes and adhesive papillae on the posterior ends. Both have rhabdites sparsely present over the entire body, concentrated at the posterior end and a preoral intestine extending to the brain. Moreover, both species inhabit the Northern Atlantic in overlapping habitats, in mud on opposite sides of the English Channel: Baie de la Forêt near Concarneau for *M. lucidum*, and the harbor near Plymouth for *M. hamatum*. Unfortunately, Furhmann provided no images of *M. lucidum* apart from a drawing of a single rhabdite bundle and was unable to collect specimens with sexual anatomy. Nevertheless, the similarities in morphology and ecology suggest it is highly likely that these represent the same species if—as we have interpreted it—the “±” of Westblad’s (1953) description means that some individuals may not have dark grey spots. Regardless, body pigmentation is known to be highly variable in this genus (Stirewalt 1937; Luther 1960), and Westblad (1953) gave no explanation on how the two species should be distinguished. In this case, *Microstomum hamatum* WESTBLAD 1953 must be considered a junior synonym for *Microstomum lucidum* FUHRMANN 1896.

### Species Descriptions

> Order Macrostomida KARLING, 1940
>
> Family Microstomidae LUTHER, 1907
>
> Genus *Microstomum* SCHMIDT, 1848

### *Microstomum afzelii* sp. nov

**Fig. 3:**
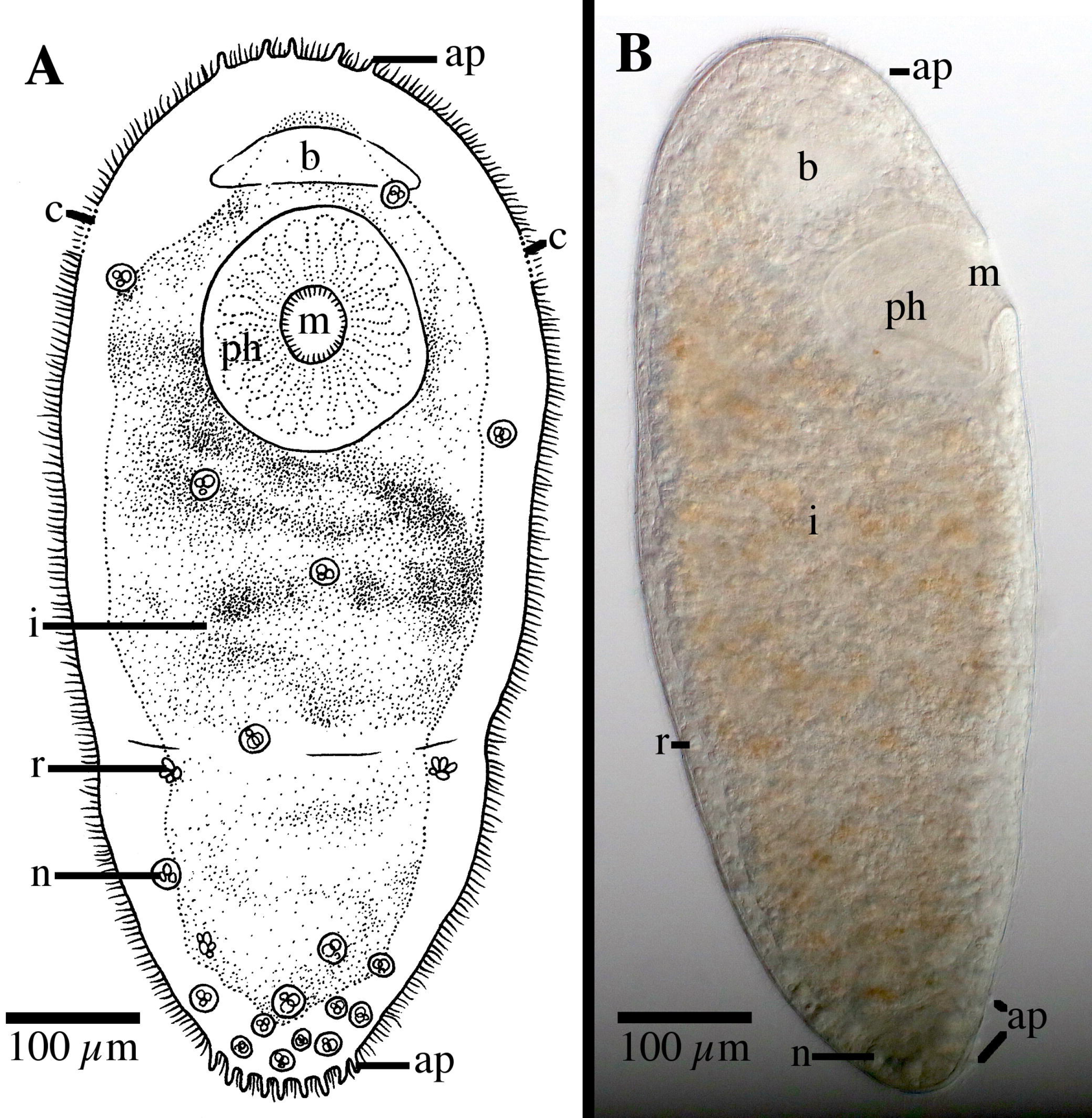
*Microstomum afzelii* sp. nov. A. Drawing of ventral view. B. Micropictograph of side view. ap adhesive papillae; b brain; c ciliary pits; i intestine; m mouth; n nematocysts; ph pharynx; r rhabdites

#### Diagnosis

Marine *Microstomum* with an almost elliptical body shape; body length 0.8 mm, 2 zooids. Colorless and without pigmented eyespots. Anterior end smoothly rounded. Posterior end slightly more tapered but also with rounded tip. Ciliary pits small and shallow; may be difficult to discern in some specimens. Few adhesive papillae present on anterior and posterior ends of body. Rhabdites sparsely present throughout body. Nematocysts particularly concentrated at posterior end. Preoral intestine extending anterior to brain. Reproductive system unknown. GenBank accession number for partial CO1 sequence XXX; for 18S XXX.

#### Etymology

Named for Lars Fredrik Afzelius, the first director of the Tjärnö Marine Laboratory, in appreciation of his contributions to the marine scientific community.

#### Material examined

Holotype: asexual specimen, Sweden, Stömstad, Saltö, 58°52’29’’N 11°08’41’’ E; 18 June 2016; 10 cm depth, marine, eulittoral sand, S. Atherton leg. (SMNH-Type-XXX; GenBank Accession CO1 XXX, 18S XXX)

Additional materials: asexual specimen; same as type (SMNH-XXX; GenBank Accession CO1 XXX, 18S XXX)

#### Description

*Microstomum* with a total body length of 0.8 mm, 2 zooids. Body colorless and without pigmented eyespots. Body shape almost elliptical, with widest point occurring approximately at midpoint or slightly anterior, lightly decreasing in width to a rounded anterior end, and a slightly more pronounced tapering to the rounded posterior end. Ration of body width:length ~1:2.5 in slightly compressed animal.

Ciliary pits weakly present, may be difficult to distinguish in some specimens. Small and shallow, located at level of posterior brain.

Epidermis uniformly covered with cilia. A few longer sensory cilia present on posterior end. Nematocysts present, highly concentrated at posterior. Rhabdites present, 15-20 μl long, sparsely scattered across entire body. Approximately 6-8 adhesive papillae, 10-15 μm long, on the anterior end; a further 10-12 located at the posterior rim.

Mouth circular, 50 μm in diameter, surrounded by cilia. Pharynx also ciliated, 138 μm long, surrounded by relatively few pharyngeal glands. Preoral intestine extending anterior to brain. Intestine filled with lightly orange-brown food.

Reproductive system unknown.

#### Notes

There are currently 23 species of *Microstomum* that have neither pigmented eyespots nor distinct body pigmentation. Of these, *M. afzelii* sp. nov. can generally be easily separated by its body shape, especially the rounded body ends, preoral intestine extending anterior to the brain and the presence of the adhesive papillae on the anterior end. The other species of *Microstomum* without very distinct and characteristic pigmentation can specifically be separated from *M. afzelii* sp. nov. in the following ways:

*M. album* (ATTEMS 1896) – anterior proboscis present; more numerous posterior adhesive papillae, presence of adhesive papillae on midbody; more numerous rhabdites; short preoral intestine
*M. bispiralis* STIREWALT 1937 – posterior end pointed; preoral intestine short; adhesive papillae and rhabdites absent; limnic habitat; see also Figs. 1, 2
*M. breviceps* MARCUS 1951 – very distinct spatulate posterior end; biparte pharynx
*M. canum* (FUHRMANN 1894) – longer, more pointed anterior end; posterior end pointed; ciliary pits deep; adhesive papillae and rhabdites absent; limnic habitat
*M. crildensis* FAUBEL 1984– short preoral intestine; anterior adhesive papillae absent; rhabdites absent; distinct constriction of body width at level of brain; see also Figs. 1, 2
*M. davenporti* GRAFF 1911– anterior adhesive papillae absent; more numerous anterior rhabdites; biparte pharynx
*M. edmondi* ATHERTON & JONDELIUS 2018 – posterior end blunt; deep bottle-shaped ciliary pits; anterior adhesive papillae absent; more numerous rhabdites; see also Figs. 1, 2
*M. giganteum* HALLEZ 1878 – very large body size; preoral intestine short; adhesive papillae and rhabdites absent; limnic habitat
*M. inexcultus* sp. nov. – rhabdites absent; anterior adhesive papillae absent; more numerous posterior adhesive papillae; see also Figs. 1, 2
*M. jenseni* RIEDEL, 1932 – posterior end tail-like; preoral intestine only reaching to level of brain; anterior adhesive papillae absent
*M. lineare* (MÜLLER 1773) – often with anterior red eyestripes; posterior end tail-like; adhesive papillae appearing absent or very small; rhabdites absent; limnic or brackish water habitat; see also Figs. 1, 2
*M. lotti* sp. nov. – distinct spatulate tail-plate; very large ciliated pits; anterior crown of adhesive papillae; small preoral intestine; see also Figs. 1, 2
*M. lucidum* (FUHRMANN 1896) – preoral intestine extending only to level of brain; anterior adhesive papillae absent
*M. marisrubri* sp. nov. – rhabdites absent; more numerous posterior adhesive papillae; anterior end longer; preoral intestine small; see also Figs. 1, 2
*M. mundum* GRAFF 1905 – characteristic lateral bulges in intestine; preoral intestine extending only to level of brain; posterior end tail-like; anterior adhesive papillae absent
*M. papillosum* (GRAFF 1882) – more numerous rhabdites at anterior end; adhesive papillae scattered across entire body; see also Figs. 1, 2
*M. rhabdotum* MARCUS 1950 – biparte pharynx; paired stripes of rhabdite bundles at ciliary pits; preoral intestine extending only to level of brain
*M. spiculifer* FAUBEL 1974 – anterior adhesive papillae absent; more numerous posterior rhabdites; longer preoral intestine reaching almost to very tip of body.
*M. trichotum* MARCUS 1950 – long and tapered anterior end; more numerous rhabdites concentrated at anterior; anterior adhesive papillae absent; preoral intestine extending only to level of brain
*M. ulum* MARCUS 1950 – preoral intestine short; deep ciliary pits; distinct spatulate tail plate; more numerous rhabdites concentrated at posterior end
*M. weberi* sp. nov. – bottle-shaped ciliary pits; anterior crown of adhesive papillae; see also Figs. 1, 2
*M. westbladi* nom. nov. – more numerous rhabdites highly concentrated at anterior end; adhesive papillae scattered across entire body; characteristic shape of field of adhesive papillae at anterior end; see also Figs. 1, 2
*M. zicklerorum* ATHERTON & JONDELIUS 2018 – individuals may have anterior red eyestripes; posterior end tail-like; adhesive papillae and rhabdites absent; limnic habitat; see also Figs. 1, 2

### *Microstomum curinigalletti* sp. nov

**Fig. 4:**
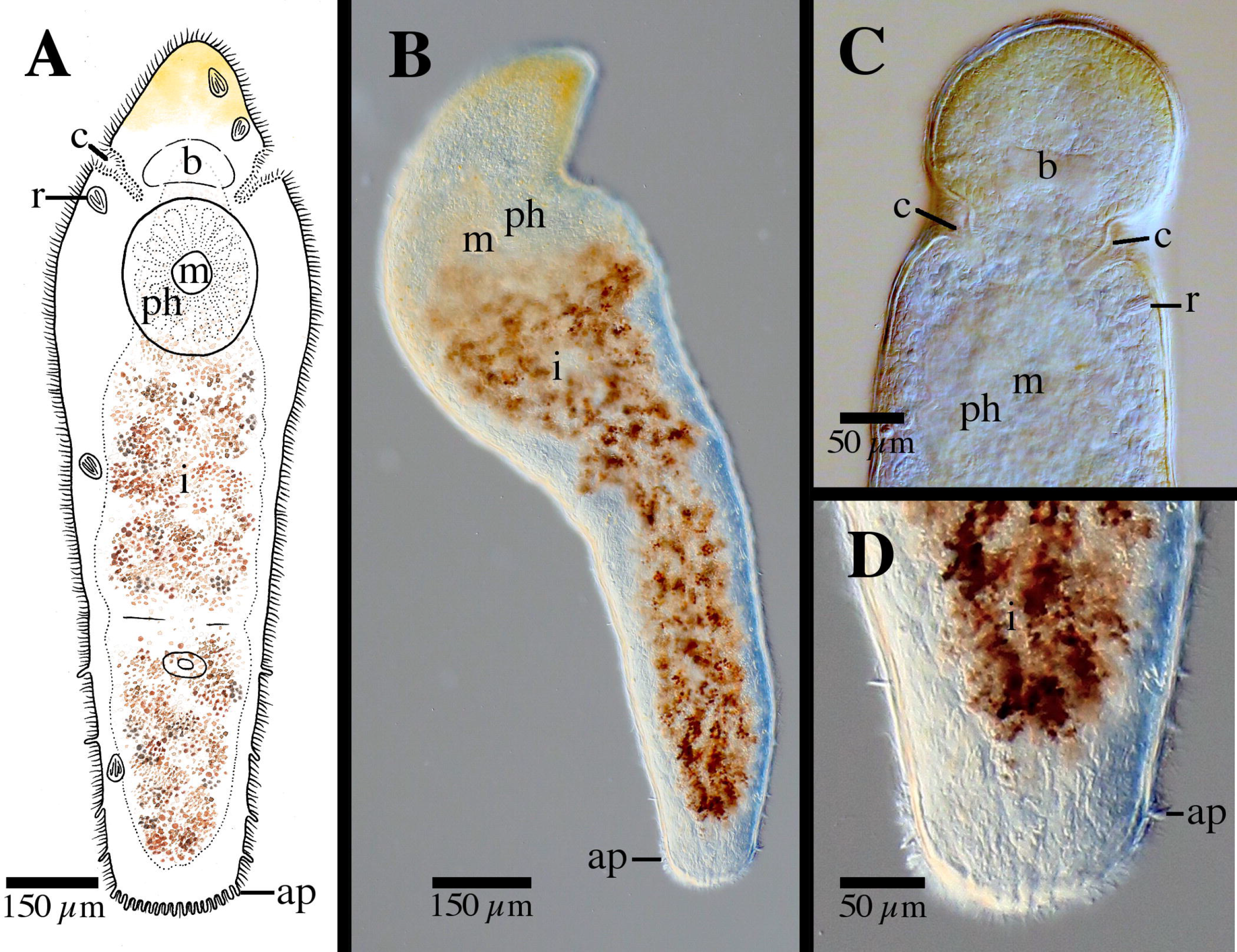
*Microstomum curinigalletti* sp. nov. A. Drawing of whole body. B. Micropictograph of whole body C. Anterior end with focus on the ciliary pits. D. Posterior end. ap adhesive papillae; b brain; c ciliary pits; i intestine; m mouth; ph pharynx; r rhabdites

#### Diagnosis

Marine *Microstomum* with a body length of 1.4 mm, 2 zooids. Anterior body end colored ochre to dark yellow-orange. Body otherwise colorless without pigmented eyespots. Body shape with a conically rounded anterior end and wide, blunt posterior end. Numerous adhesive papillae present at posterior end. Rhabdites present but few, scattered across the body. Ciliary pits large and bottle-shaped, located anterior to pharynx. Preoral intestine short. Reproductive system unknown. GenBank accession number for partial CO1 sequence XXXX; for 18S XXX.

#### Etymology

This species is dedicated with many thanks to Dr. Marco Curini-Galletti, who found the first specimens of these animals.

#### Material examined

Holotype: Asexual specimen; Sweden, Fiskebäckskil, Kristineberg Sven Lovén Center for Marine Research, 58°14′59′′ N, 11°26′45′′ E, 20 Aug. 2015, marine, sublittoral phytal on algae, M. Curini-Galletti leg. (SMNH-Type-XXXX; GenBank Accession 18S XXX)

Additional materials: 5 asexual; same as holotype (SMNH-XXXX; GenBank Accession 18S XXX)

#### Description

*Microstomum* with a total body length to 1.4 mm, 2 zooids. Anterior body end colored ochre to dark yellow-orange. Body otherwise colorless and reflecting intestines. Pigmented eyespots absent. Body shape with a conically rounded anterior end; posterior end blunt and almost as wide as rest of body. Ratio of width generally consistent across body, width:length is 1:4 in slightly compressed animal.

Ciliary pits large, clearly distinguishable anterior to pharynx; bottle shaped, extending at a 45° angle posterior into the body ~65 μm; opening 30 μm long.

Epidermis uniformly covered with cilia. Nematocysts present. Few rhabdites, to 25 μm long, scattered throughout the body. Numerous adhesive papillae, 12-15 μm long, present at posterior tip.

Mouth small and circular surrounded by cilia. Pharynx surrounded by normal ring of pharyngeal glands. Preoral intestine very short, hardly extending past mouth.

Reproductive system unknown

#### Notes

In body shape and size and, especially, size, shape and location of ciliary pits (Fig. 4C), *Microstomum curinigalletti* sp. nov. is most similar to *M. edmondi*. The two species also share habitats and distributions and were collected in the same samples. *M. curinigalletti* sp. nov., however, can easily be distinguished from *M. edmondi* through the yellow pigmentation of the anterior end and the shorter preoral intestine. Results from the phylogenetic and bPTP analyses also clearly separated the two species (Figs. 1, 2).

### *Microstomum inexcultus* sp. nov

**Fig. 5:**
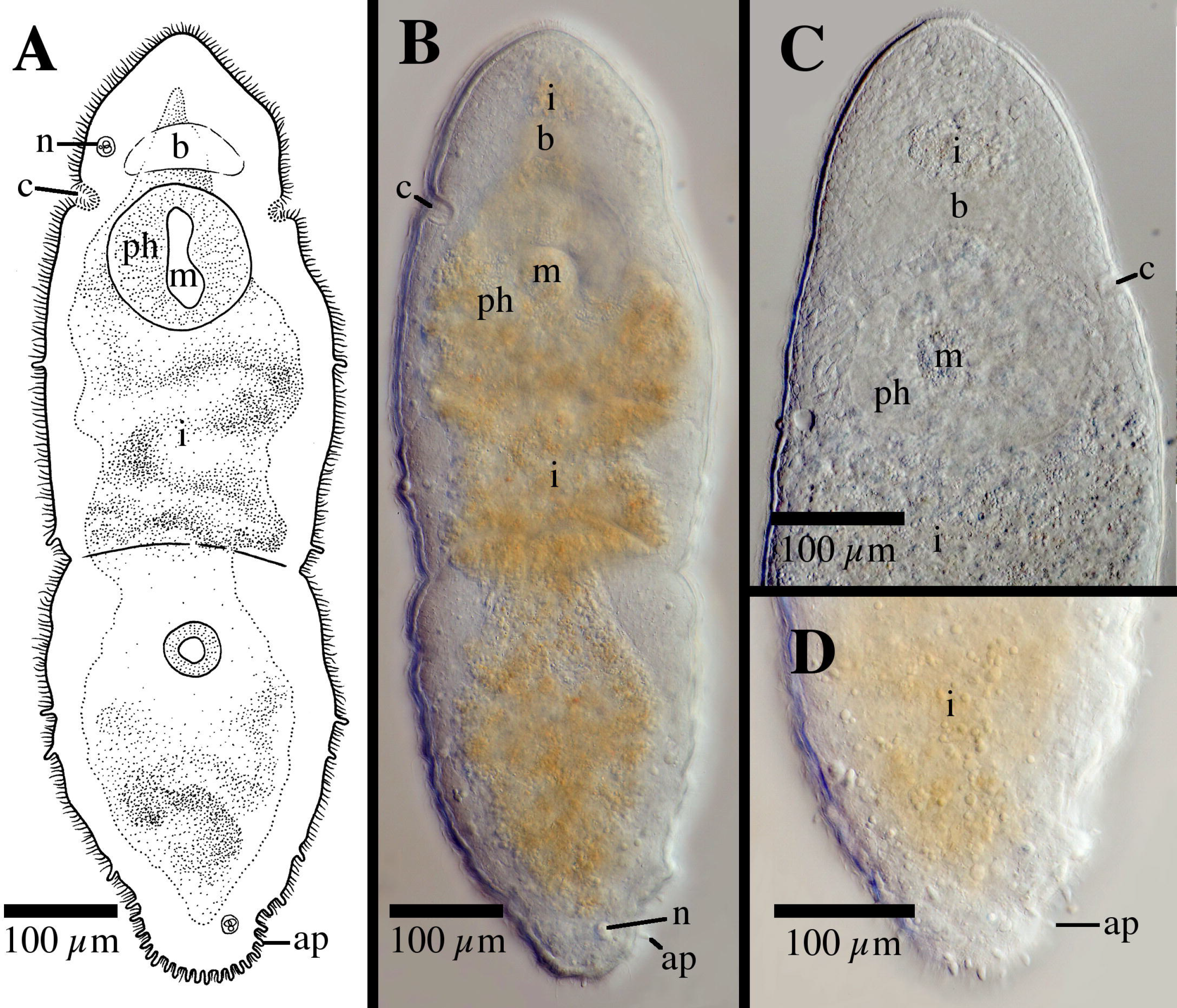
*Microstomum inexcultus* sp. nov. A. Drawing of whole body. B. Micropictograph of whole body C. Anterior end. D. Posterior end with focus on adhesive papillae. ap adhesive papillae; b brain; c ciliary pits; i intestine; m mouth; n nematocysts; ph pharynx

#### Diagnosis

Marine *Microstomum* with a body length of 0.85, 2 zooids. Colorless and without pigmented eyespots. Body shape with a conically pointed anterior end and a tapered posterior end with a blunt tip. Numerous adhesive papillae present at posterior end. Rhabdites absent. Ciliary pits clearly present. Preoral intestine long, stretching almost to anterior body end. Reproductive system unknown. GenBank accession number for partial CO1 sequence XXXX; for 18S XXX.

#### Etymology

This species named for its very simple appearance without rhabdites (*inexcultus*, Latin: unadorned, unembellished)

#### Material examined

Holotype: Asexual specimen; Sweden, Stömstad, Saltö, 58°52’29’’N 11°08’41’’ E; 18 Jun. 2016; 10 cm depth, marine, eulittoral sand, S. Atherton leg. (SMNH-Type-XXXX; GenBank Accession CO1 XXX, 18S XXX)

Additional materials: 5 asexual specimens, same as holotype; 2 asexual specimens Sweden, 58°25’24’’N 11°39’34’’ E; 15 Jun. 2015; 1 m depth, marine; S. Atherton leg. (SMNH-XXXX; GenBank Accession CO1 XXX, 18S XXX)

#### Description

*Microstomum* with a total body length to 0.85 mm, 2 zooids. Body colorless and without pigmented eyespots. Body shape with a conically pointed anterior end and a tapered posterior end with a blunt tip. Maximum body width just posterior to pharynx; ratio of maximum width:length is ~1:3 in slightly compressed animal.

Ciliary pits clearly distinguishable at level of anterior pharynx 170 μm from anterior body end; roundish to cup-shaped, extending at a slight angle into the body 27 μm; opening 20 μl long.

Epidermis uniformly covered with cilia. Very few nematocysts present, primarily at body ends. Rhabdites absent. Numerous adhesive papillae, ~15 long, present at posterior end and along up to ~⅕ of the posterior body; may also be present at zone of division in very well developed zooids.

Mouth elliptical, 45 μm long, surrounded by cilia. Pharynx 125 μm long, surrounded by normal ring of pharyngeal glands. Preoral intestine extending well anterior to brain, reaching almost to anterior body tip.

Reproductive system unknown

#### Notes

*Microstomum inexcultus* sp. nov. can be distinguished from all other species of *Microstomum* through a combination of the following characters: absent of pigmentation or eyespots; preoral brain very long and extending almost to anterior body tip (Fig. 5C); rhabdites absent. The absence of rhabdites in particular distinguished this species, since only 11 other species of *Microstomum* do not have rhabdites recorded; these can be distinguished from *M. inexcultus* sp. nov. by the following:

*M. artoisi* ATHERTON & JONDELIUS 2018 – anterior red eyestripes present; preoral intestine short; posterior end tail-like; adhesive papillae absent; limnic habitat; see also Figs. 1, 2
*M. bispiralis* STIREWALT 2018 – preoral intestine short; posterior end pointed; adhesive papillae absent; limnic habitat; see also Figs. 1, 2
*M. caerulescens* (SCHMARDA, 1859) – body blue; adhesive papillae absent; limnic habitat
*M. canum* (FUHRMANN 1894) – posterior end tail-like; ciliary pits deep; adhesive papillae absent; limnic habitat
*M. crildensis* FAUBEL, 1984 – more rounded anterior; constriction of body width at level of brain; short preoral intestine; see also Figs. 1, 2.
*M. giganteum* HALLEZ 1878 – very large body size; preoral intestine short; adhesive papillae absent; limnic habitat
*M. lineare* (MÜLLER, 1773) – anterior red eyestripes often present; posterior end tail-like; adhesive papillae appearing very small or absent; limnic or brackish water habitat; see also Figs. 1, 2
*M. marisrubri* sp. nov. – longer frontal end; small preoral intestine; anterior adhesive papillae present; see also Figs. 1, 2.
*M. punctatum* DORNER 1902 – body brown-yellow with black spots; preoral intestine short; posterior end tail-like; limnic habitat
*M. tchaikovskyi* ATHERTON & JONDELIUS 2018 – anterior red eyestripes present; preoral intestine short; posterior end tail-like; adhesive papillae absent; limnic habitat; see also Figs. 1, 2
*M. zicklerorum* ATHERTON & JONDELIUS 2018 – anterior red eyestripes may be present; preoral intestine short; posterior end tail-like; adhesive papillae absent; limnic habitat; see also Figs. 1, 2

### *Microstomum lotti* sp. nov

**Fig. 6:**
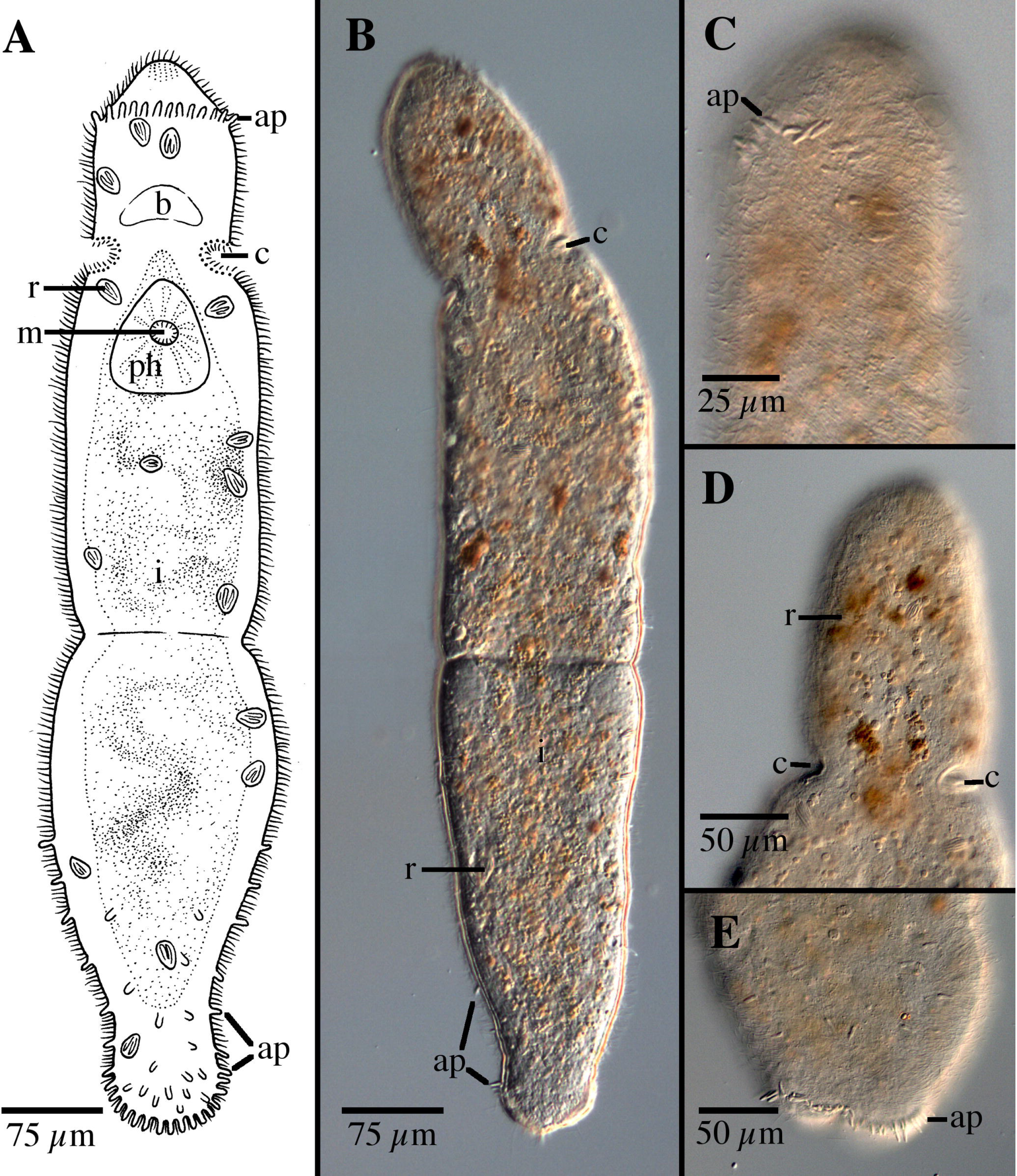
*Microstomum lotti* sp. nov. A. Drawing of whole body. B. Micropictograph of whole body C. Anterior end with focus on adhesive papillae. D. Anterior end with focus on ciliary pits E. Posterior end with focus on adhesive papillae. ap adhesive papillae; b brain; c ciliary pits; i intestine; m mouth; ph pharynx; r rhabdites

#### Diagnosis

Marine *Microstomum* with a body length of 0.8, 2 zooid. Body colorless without pigmented eyespots. Anterior end conical; posterior end forming a spatulate tail-plate set off by a constriction. Adhesive papillae present along posterior half of the body and very numerous at tail plate. An additional row of adhesive papillae present ventrally at the frontal region ~30 μm from anterior tip. Ciliary pits rounded. Preoral intestine short. Rhabdites present. Reproductive system unknown. GenBank accession for 18S KP730483, KP730493.

#### Etymology

This species is named in honor of Christian Lott, who was a co-organizer of the 2010 BIOSAND workshop, where the sample was collected.

#### Material examined

Holotype: Asexual specimen; Italy, Sant Andrea Bay, 42°48′31′′ N, 10°08′30′′ E, 26 April 2010, marine (MTP LS 660; GenBank Accession 18S KP730483) Other materials: same as holotype (MTP LS 635; GenBank Accession 18S KP730493)

#### Description

*Microstomum* with a total body length to 0.8 mm, 2 zooids. Body colorless and without pigmented eyespots. Anterior end conically pointed with row of adhesive papillae at the frontal end, ~30 μm from anterior tip. Posterior end forming a distinct tail-plate separated by a constriction of the body width. Width to length ration is approximately 1:5.

Ciliary pits large, present posterior to brain; each forming a roundish indentation angled slightly anteriorly; extending inward 21 μm with 16 μm long openings.

Epidermis uniformly covered with cilia. Rhabdites bundles scattered throughout the body, ~12-17 μm. In addition to those at the frontal end, numerous adhesive papillae, 10 μm long, are present at the tail-plate and less frequently at the margins to approximately halfway up the body.

Mouth circular and surrounded by cilia. Pharynx 220 μm long, surrounded by normal ring of pharyngeal glands. Preoral intestine reaching to level halfway between pharynx and brain.

Reproductive system unknown.

#### Notes

The only species of *Microstomum* with distinct spatulate tail-plates are *M. breviceps* and *M. ulum. M. lotti* sp. nov. differs from *M. breviceps* by the row of adhesive papillae on the frontal end (Fig. 8C) and by the small preoral intestine. *M. lotti* sp. nov. is more similar to *M. ulum*, although the two species still can be differentiated by the field of rhabdites at the frontal end of *M. ulum* and the row of adhesive papillae at the frontal end of *M. lotti* sp. nov.

### *Microstomum marisrubri* sp. nov

**Fig. 7:**
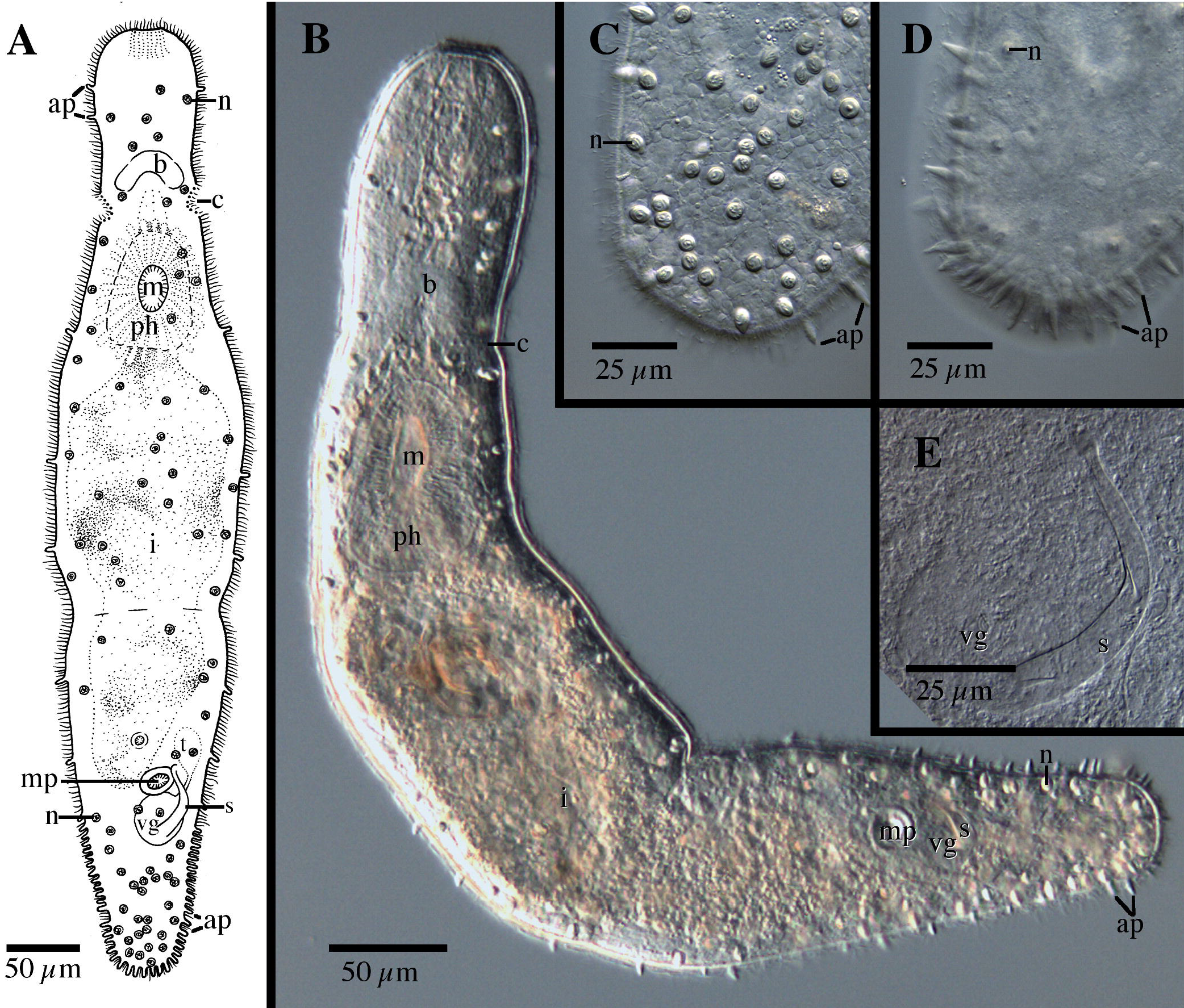
*Microstomum marisrubri* sp. nov. A. Drawing of whole body. B. Micropictograph of whole body C. Posterior end with focus on the nematocysts. D. Posterior end with focus on the adhesive papillae. E. Male stylet. ap adhesive papillae; b brain; c ciliary pits; i intestine; m mouth; mp male pore; n nematocyst; ph pharynx; r rhabdites; s male stylet; t testis; vg vesicula granulorum

#### Diagnosis

Marine *Microstomum* with a body length of 0.65, 1 zooid. Body colorless without pigmented eyespots. Anterior end large, with a rounded tip. Posterior end blunt. Adhesive papillae sparsely present over entire body but extremely numerous at posterior ⅕ of body. Ciliary pits present, cup shaped, located at level between anterior pharynx and brain. Preoral intestine short, reaching just anterior to pharynx. Rhabdites absent, but with many nematocysts intensely concentrated at posterior end. Male stylet a curved tube tapering distally to a slightly enlarged end. GenBank accession number for partial CO1 sequence KP730580; for 18S KP730505.

#### Etymology

This species is named after the collection location *(maris*, Latin: sea; *rubri* Latin: red)

#### Material examined

Holotype: ◻; Egypt, Mangrove Bay, 25°52′15′′ N, 34°25′04′′ E, 11 January 2009, marine (lactophenol preparation NHMUK-2015.1.30.22; digital morphological voucher MTP LS 524; GenBank Accession CO1 KP730580, 18S KP730505)

#### Description

*Microstomum* with a total body length to 0.65 mm, 2 zooids. Body colorless and without pigmented eyespots. Anterior end long, measuring 95 μm from tip to anterior brain, with a widely rounded tip. Body width slightly decreasing from frontal end to ciliary pits, slightly increasing toward the middle of the animal and then decreasing again to a blunt posterior end. Animal measures approximately 80:70:120:45 μm at frontal end: ciliary pits: body middle: posterior end.

Ciliary pits present posterior to brain; each forming a roundish indentation angled at approximately 45° into the body; extending 10 μm with 10 μm long openings.

Epidermis uniformly covered with cilia. Rhabdites absent. Very many nematocysts present throughout the animal and especially intensely concentrated at the posterior end. Numerous adhesive papillae, ~10 μm long, present along ~⅕ of the posterior body and also sparsely occurring anteriorly.

Mouth surrounded by cilia. Pharynx 95 μm long, surrounded by normal ring of pharyngeal glands. Preoral intestine short, terminating just posterior to level of brain.

Male reproductive system with single developing testis connected by short vas deferens to a rounded vesicula granulorum. Male stylet, 63 μm long, a slightly tapering tube, 9 μm wide at base and ~3 μm at narrowest part of distal end; curving a little less than 90°, with a slight widening at the distal tip to ~4.5 μm. Genital pores separate. Male pore 18 μm in diameter.

#### Notes

The stylet and reproductive system of *Microstomum marisrubri* sp. nov. (Fig. 7E) is very similar to *M. papillosum*, although it is smaller (62 μm compared to 80 μm, respectively). Other differences between the two species can be found in the size of the preoral intestine, which *for M. papillsum* extends past the brain, and the presence of numerous rhabdites at the anterior end. As discussed above, few species of *Microstomum* lack rhabdites, and these can be distinguished:

*M. artoisi* ATHERTON & JONDELIUS 2018 – anterior red eyestripes present; posterior end tail-like; adhesive papillae absent; limnic habitat; see also Figs. 1, 2
*M. bispiralis* STIREWALT, 1937 – posterior body pointed; adhesive papillae absent; limnic habitat; see also Figs. 1, 2.
*M. caerulescens* (SCHMARDA, 1859) – body blue; preoral intestine extending past brain; adhesive papillae absent; limnic habitat
*M. canum* (FUHRMANN 1894) – posterior end tail-like; adhesive papillae absent; limnic habitat
*M. crildensis* FAUBEL, 1984 – shape and size of the male stylet; lack of anterior adhesive papillae and fewer posterior adhesive papillae; see also Figs. 1, 2.
*M. giganteum* HALLEZ 1878 – very large body size; posterior end a thin tail; adhesive papillae absent; limnic habitat
*M. inexcultus* sp. nov. – smaller frontal end; large preoral intestine; lack of anterior adhesive papillae; see also Figs. 1, 2.
*M. lineare* (MÜLLER, 1773) – anterior red eyestripes often present; posterior end tail-like; adhesive papillae very small or appearing absent; limnic or brackish water habitat; see also Figs. 1, 2
*M. punctatum* DORNER 1902 – body brown-yellow with black spots; posterior end tail-like; limnic habitat
*M. tchaikovskyi* ATHERTON & JONDELIUS 2018 – anterior red eyestripes present; posterior end tail-like; adhesive papillae absent; limnic habitat; see also Figs. 1, 2
*M. zicklerorum* ATHERTON & JONDELIUS 2018 – anterior red eyestripes may be present; posterior end tail-like; adhesive papillae absent; limnic habitat; see also Figs. 1, 2

### *Microstomum schultei* sp. nov

**Fig. 8:**
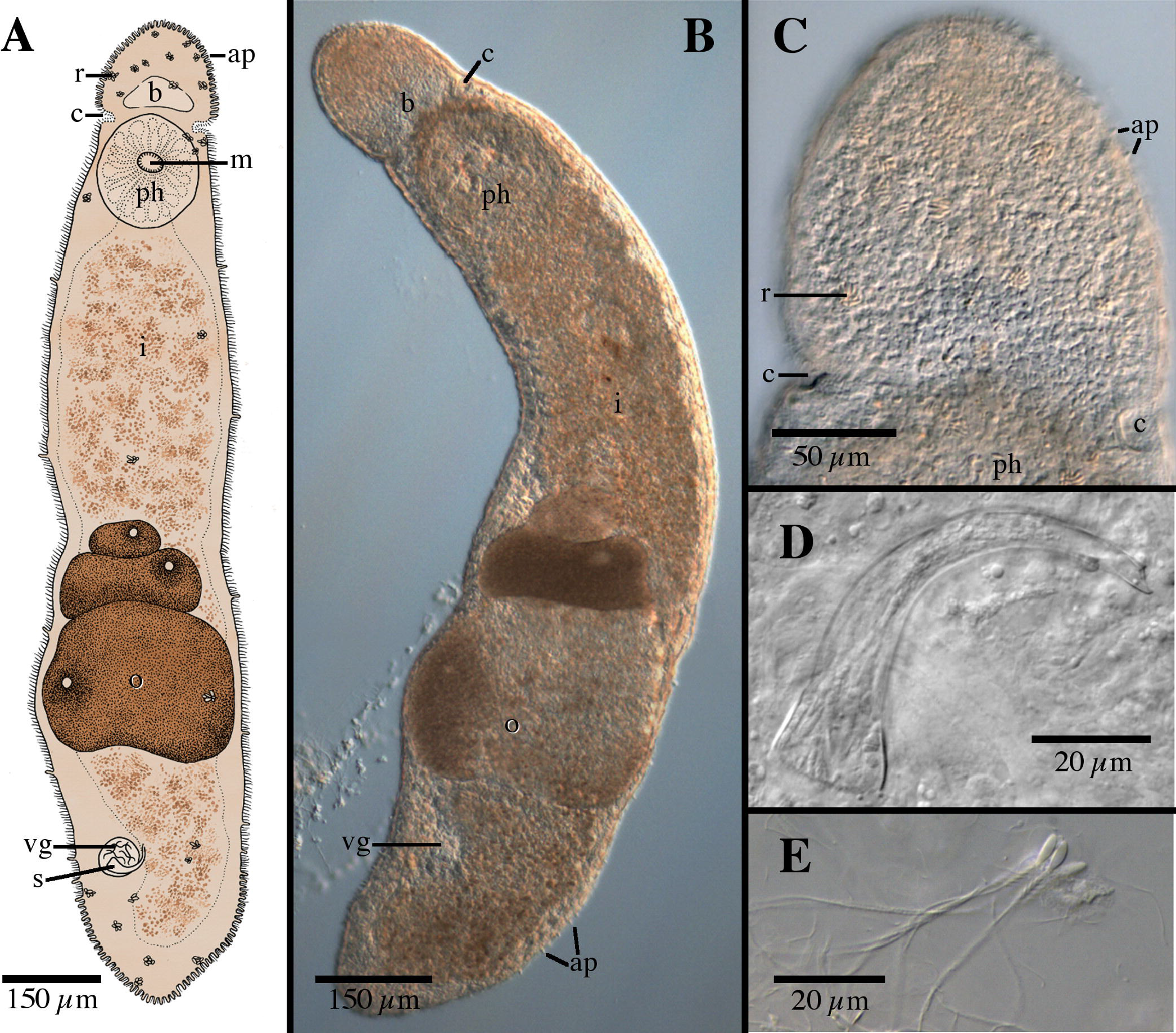
*Microstomum schultei* sp. nov. A. Drawing of whole body. B. Micropictograph of whole body C. Anterior end with focus on the ciliary pits. D. Male stylet. E. Sperm. ap adhesive papillae; b brain; c ciliary pits; i intestine; m mouth; o ovary; ph pharynx; r rhabdites; s male stylet; vg vesicula granulorum

#### Diagnosis

Marine *Microstomum* with a body length of 1.5 mm, 1 zooid. Body appears light reddish-brown without pigmented eyespots. Anterior end rounded and set apart from the rest of the body by a slight constriction at the ciliary pits. Posterior end conically blunted. Adhesive papillae and rhabdites concentrated at anterior and posterior ends. Ciliary pits present, cup shaped. Preoral intestine short. Reproductive system with curved male stylet, tapering from a wider base to a narrow tube with a beveled tip. GenBank accession number for 18S KP730494.

#### Etymology

This species is named in honor of Gregor Schulte, who was a field assistant during the 2010 BIOSAND workshop and aided in sample collection.

#### Material examined

Holotype: ◻; Italy, Fetovaia Bay, 42°43′36′′ N, 10°09′33′′ E, 05 May 2010, marine (lactophenol preparation NHMUK-2015.1.30.21; digital morphological voucher MTP LS 700; GenBank Accession 18S KP730494)

#### Description

Microstomum with a total body length to 1.5 mm, 1 zooid. Body light reddish-brown, without pigmented eyespots. Anterior end rounded and set apart from the rest of the body by a slight constriction at the ciliary pits. Posterior end conically blunted. Ratio of maximum width:length is ~1:5 in slightly compressed animal.

Ciliary pits present at level just posterior to brain, 170 μm from anterior body end; roundish to cup-shaped, extending into the body 30 μm with 15 μm long openings.

Epidermis uniformly covered with cilia. Rhabdites present throughout the body, but especially concentrated at the anterior and posterior ends. Numerous short adhesive papillae, ~10 μm long, present, heavily concentrated at posterior end along ~⅙ of the body and also anterior to brain.

Mouth circular, 40 μm long, surrounded by cilia. Pharynx 185 μm long, surrounded by normal ring of pharyngeal glands. Preoral intestine short.

Reproductive with single medial ovary and caudally developing eggs. Male system with vesicula granulorum connected to male stylet, 60 μm long, curved and tapering from a wider base to form a narrow tube with a beveled tip. Stylet base 20 μm wide, distal tube ~4 μm wide, opening ~5 μm long. Sperm ~70 μm long, with a rounded head and a single flagellum with lateral setae.

#### Notes

There are a few species of *Microstomum* with a similarly shaped male stylet (Fig. 6D). *M. schultei* sp. nov. can be separated from each of these through:

*M. crildensis* FAUBEL, 1984 – stylet much smaller (32 μm instead of 60 μm), with a more narrow base; rhabdites absent; anterior adhesive papillae absent; body colorless; see also Figs. 1, 2
*M. gabriellae* MARCUS, 1950 – stylet even more curved, tracing an entire half circle 180°; terminal end not beveled; preoral intestine large, extending past brain; red eyespot present
*M. septentrionale* (SABUSSOW 1900)– stylet without beveled terminal end; preoral intestine reaching to level of anterior brain; body color with dorsal yellow pigmentation; see also Figs. 1, 2.

In addition, *M. schultei* sp. nov can generally be distinguished from the other species of *Microstomum* through a combination of the presence of adhesive papillae at the anterior and posterior ends and a short preoral intestine (defined here as a preoral intestine that does not extend to at least the level of posterior brain; Fig. 6). *M. schultei* sp. nov. can be separated from the other species of *Microstomum* with these characters by the following ways:

*M. album* (ATTEMS 1896) – body colorless or white; anterior proboscis present; ciliary pits small
*M. crildensis* FAUBEL, 1984 – see above
*M. curinigalletti* sp. nov. – frontal end colored ochre; ciliary pits large and bottle shaped; anterior adhesive papillae absent; fewer rhabdites; see also Figs. 1, 2
*M. lotti* sp. nov. – posterior end spatulate; anterior crown of adhesive papillae; body colorless; see also Figs. 1, 2
*M. marisrubri* sp. nov. – longer frontal end; more numerous posterior adhesive papillae; rhabdites absent; body colorless; see also Figs. 1, 2
*M. punctatum* DORNER 1902 – body brown-yellow with black spots; pointed anterior and tail-like posterior ends; ciliary pits small and shallow; rhabdites absent; limnic habitat
*M. rubromaculatum* (GRAFF, 1882) - red pigmentation stripe at anterior end present; widely blunt posterior end; see also Figs. 1, 2
*M. ulum* MARCUS 1950 – pointed anterior end; distinct spatulate tail-plate at posterior end; more rhabdites concentrated at anterior end; body colorless

### *Microstomum weberi* sp. nov

**Fig. 9:**
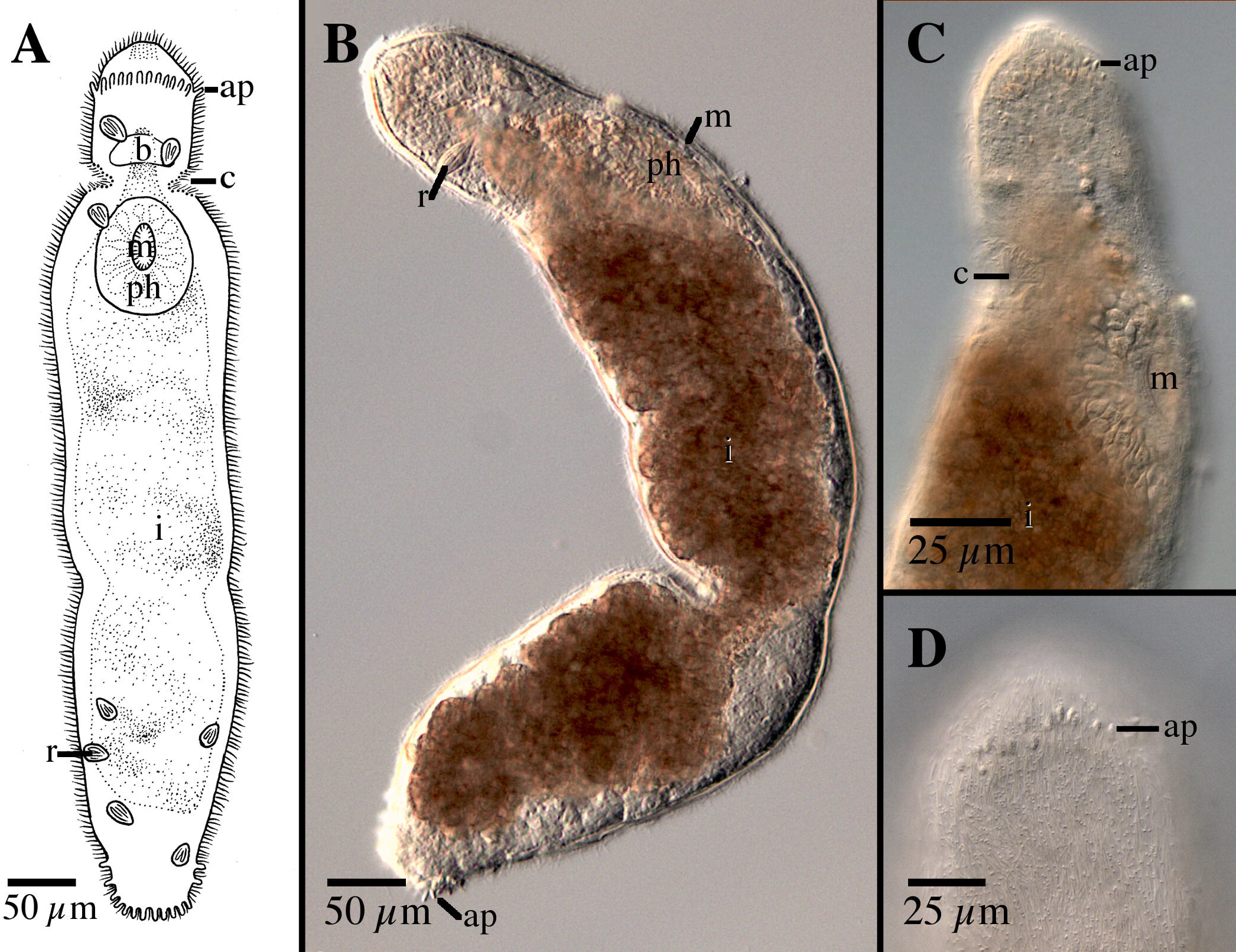
*Microstomum weberi* sp. nov. A. Drawing of whole body. B. Micropictograph of whole body C. Anterior end side view. D. Anterior end with focus on adhesive papillae. ap adhesive papillae; b brain; c ciliary pits; i intestine; m mouth; ph pharynx; r rhabdites

#### Diagnosis

Marine *Microstomum* with a body length of 0.65 mm, 2 zooids. Body colorless without pigmented eyespots. Anterior end conical; posterior end blunt. Adhesive papillae occur at posterior end and in a ventral row at the frontal end. Ciliary pits bottle-shaped. Preoral intestine extending to anterior brain or just beyond. Large rhabdites bundles present at anterior and posterior ends. Reproductive system unknown. GenBank accession number for partial CO1 sequence KP730576; for 18S KP730487.

#### Etymology

This species is named in honor of Miriam Weber, who collected the sample and was co-organizer of the 2010 BIOSAND workshop

#### Material examined

Holotype: Asexual specimen; Italy, Pianoso, 42°34′29′′ N, 10°03′59′′ E, 30 April 2010, marine (MTP LS 666; GenBank Accession CO1 KP730576, 18S KP730487)

#### Description

Body length 0.65 mm, 2 zooids. Body colorless and without pigmented eyespots. Anterior end conically rounded. Body width smaller at anterior end and increases perceptively just after ciliary pits. Posterior tapering to a wide blunt end.

Ciliary pits bottle-shaped, present at level of anterior pharynx; extending 13 μm into the body at slight posteriorly-directed angle; with 12 μm long openings.

Epidermis uniformly covered with cilia. Large rhabdites bundles, to 20 μm, present only at anterior and posterior ends. A row of adhesive papillae at the frontal end, situated ventrally 25 μm from anterior tip. Additional adhesive papillae, ~7-10 μm long, also present at posterior end, but are otherwise absent from the body margins.

Mouth eliptical and surrounded by cilia. Pharynx 105 μm long, surrounded by normal ring of pharyngeal glands. Preoral intestine reaching just past anterior brain.

Reproductive system unknown.

#### Notes

This is only the second species of *Microstomum* with a ventral row of adhesive tubules on the frontal end (Fig. 9C), the other being *M. lotti* sp. nov. *M. weberi* sp. nov. can be distinguished by general body shape and size, through the bottle-shaped ciliary pits (Fig. 9C), the longer preoral intestine and by the large rhabdite bundles present only near the body ends. Results from the phylogenetic and bPTP analyses also clearly separated the two species (Figs. 1, 2).

### *Microstomum westbladi* nom. nov

**Fig. 10:**
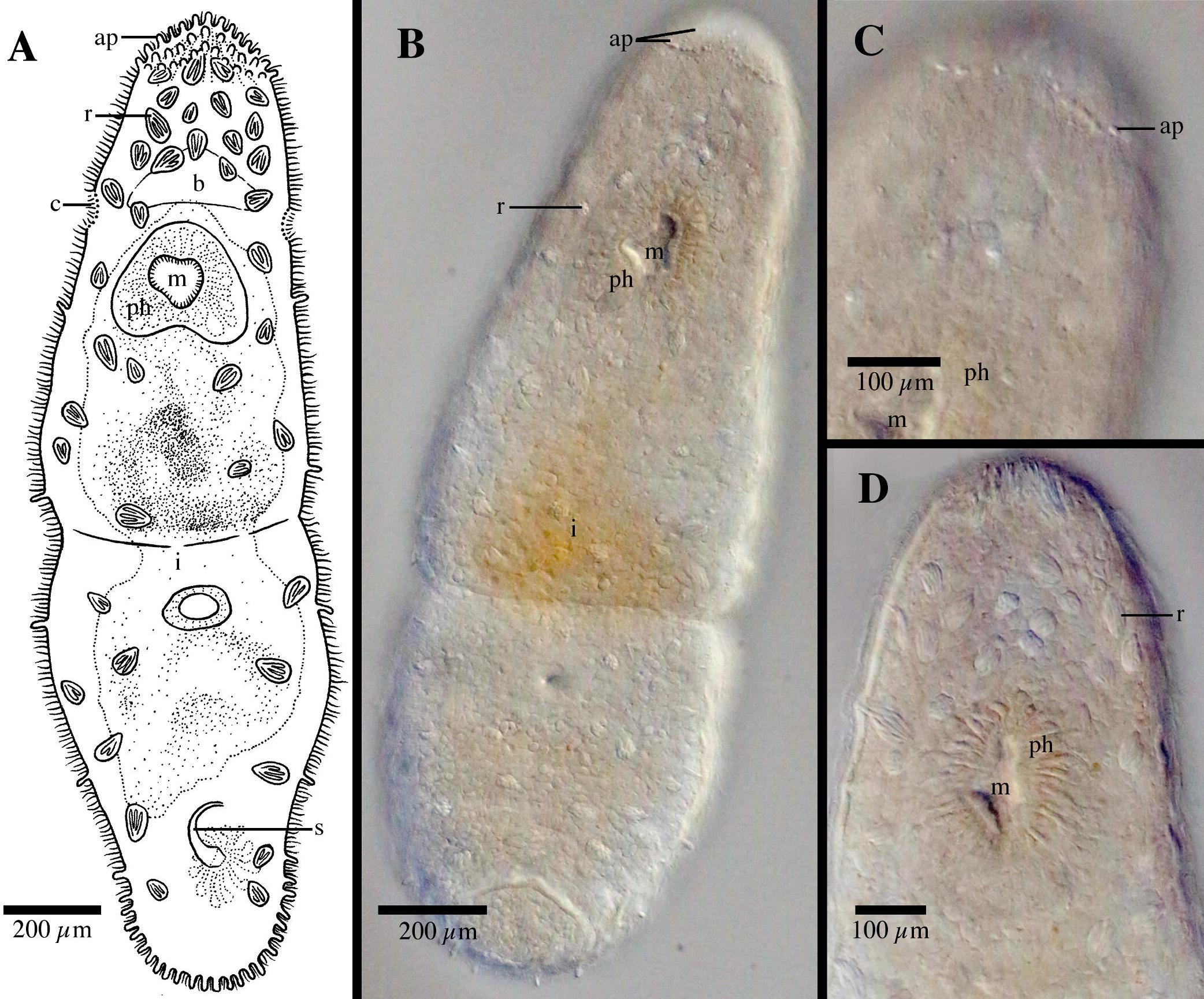
*Microstomum westbladi* nom. nov. A. Drawing of whole body. B. Micropictograph of whole body C. Anterior end focus on adhesive papillae. D. Anterior end with focus on rhabdite field. ap adhesive papillae; b brain; c ciliary pits; i intestine; m mouth; ph pharynx; r rhabdites; s male stylet

#### Synonym

*Microstomum papillosum* GRAFF 1882, sensu Westblad 1953, pg 406-7

#### Material examined

Holotype: Asexual, sectioned specimen; Norway, Herdla; 15 July 1934; 5-10 m depth, marine amphioxus sand, E. Westblad leg. (SMNH-Type-XXX74)

Additional materials: 1 asexual, sectioned, same as holotype (SMNH-94979); 1 asexual specimen, Sweden, Fiskebäckskil, 58°15’05’’N 11°27’52’’ E, 26 Aug. 2015, marine, eulittoral sand, S. Atherton leg. (SMNH-XXXX; GenBank Accession CO1 XXX, 18S XXX)

Westblad, 1953 – Norway, Herdla, Kværnepoll current; Sweden, Gullmarfjord This paper: Sweden, Fiskebäckskil, 58°15’05’’N 11°27’52’’ E

#### Diagnosis

Marine *Microstomum* with strap shaped body, to 2.0 mm long and 5 zooids. Colorless and without pigmented eyespots. Anterior end rounded. Posterior tapered to a blunt end. Ciliary pits small. Numerous rhabdites intensely concentrated at frontal end and present throughout body. Numerous adhesive papillae present at posterior end and more sparsely anterior. Characteristic grouping of adhesive papillae covering the anterior tip: field longest at both lateral sides and curving upwards toward the middle. Preoral intestine extending to brain. Reproductive system including male stylet, 60 μm long, and shaped as a smooth curve 180° curve, with a maximum width ~⅙ of the way to the distal end, decreasing proximally and also distally to a point. GenBank accession number for partial CO1 sequence XXXX; for 18S XXX.

#### Etymology

This species is named in honor of Dr. Einar Westblad, who first described these animals.

#### Materials examined

2 asexual; Sweden, Malmön, 58°20’18’’ N 11°20’26’’ E; 13 Aug. 2015; marine, 1.5 m depth; S. Atherton & Y. Jondelius leg. (SMNH-XXXX; GenBank Accession CO1 XXX, 18S XXX)

#### Description

*Microstomum* with a total body length to 2.0 mm, 5 zooids. Body colorless and without pigmented eyespots. Body strap-shaped with a bluntly rounded anterior end, an almost equal width along entire body until a slight decreasing in width to a bluntly rounded posterior end.

Ciliary pits small and shallow cups, located at level of anterior pharynx. Epidermis uniformly covered with cilia. Nematocysts not seen or recorded.

Rhabdites present, ~15-30 μm long, throughout entire body, but intensely concentrated at anterior end. Numerous adhesive papillae, ~12-15 μm long, present at posterior end and more sparsely anterior. Further adhesive papillae present at the anterior tip grouped in a very distinct pattern: the field covers the anterior tip, toupee like, i.e. longest at both lateral sides and curving upwards toward the middle.

Mouth elliptical surrounded by cilia. Pharynx 110 μm long, surrounded by normal ring of pharyngeal glands. Preoral intestine extending to brain.

Reproductive system mostly unknown. Following Westblad (1953) stylet recorded as 60 μm long, shaped as a smooth curve 180° curve, with the width first increasing along the most proximal ~⅙ of the stylet and then smoothly decreasing in width to a pointed, terminal opening.

#### Notes

*Microstomum westbladi* nom. nov. can be readily distinguished from *M. papillosum* by the characteristic pattern of the anterior adhesive papillae (Fig. 10B, C), the very high concentration of rhabdites in the anterior end (Fig. 10D) and the shape of the stylet, which decreases in width toward the proximal end as well as distally. In addition, results of the phylogenetic and bPTP analyses clearly separate the two species (Figs. 1, 2).

*M. westbladi* nom. nov. is most similar to *M. septentrionale*, particularly since both species are of similar size and shape, have a very high concentration of rhabdites in the anterior end, and are known to have overlapping habitats. However, *M. westbladi* nom. nov. can be distinguished morphologically by its colorless body without dark or orange-yellow in the anterior and through the characteristic pattern of anterior adhesive papillae. Additionally, the stylet shape of *M. westbladi* nom. nov. differs by decreasing in width toward the proximal end. *M. septentrionale* and *M. westbladi* nom. nov. are closely related sister taxa (Figs. 1, 2).

## Discussion

### Phylogenetic Analyses and DNA Taxonomy

The use of nucleotide sequences to delineate species and capture biodiversity is an important advancement in Microstomidae taxonomy. As noted in the introduction, the animals may have relatively few morphological characters on which to base descriptions, and thus identification from morphology alone may require high amounts of effort, specimens and specialized taxonomic expertise. Further, many morphological features are highly variable depending on environmental conditions (Stirewalt 1937; Bauchhenss 1971; Heitkamp 1982). Laboratory studies of *Microstomum lineare* have demonstrated that body shape and size varies based on food availability and sexual maturity; body color ranges from colorless, white, yellow, grey, reddish or brownish based on light, food type and habitat; and even amount of pigmentation in the characteristic red eyespots can range from large bright red slashes at the frontal end to non-existence depending on light intensity (Luther 1960; Bauchhenss 1971; Heitkamp 1982; Atherton & Jondelius 2018b). DNA taxonomy provides a robust, reproducible and relatively easy method of identification that can also account for the hidden biodiversity of cryptic species (Pérez-Ponce de León & Poulin 2016; Fonseca et al. 2017).

The results of the bPTP analyses on the concatenated 18S and CO1 datasets (Fig. 2) were able to reliably separate the species of Microstomidae and supported the more traditional morphological delineation. Support for four species (*M. crildensis, M. laurae, M. lotti, M. septentrionale*) was consistent but low (0.49-0.68) and is unsurprising given the overall low intraspecific genetic diversity within each *Microstomum* species. Support was moderate to high (≥0.79) for all other species.

Within the Microstomidae, our results found all limnic species (*M. artoisi, M. bispiralis, M. lineare, M. tchaikovskyi, M. zicklerorum*) grouped together in a single clade with high support (0.91), indicating a single instance of freshwater invasion from marine habitats. The transition of marine to freshwaters within a lineage is a common occurrence for many different animal taxa (Lee & Bell 1999, Davis et al. 2012) and has occurred independently at least five other times within the Platyhelminthes, including within Macrostomidae (Schockaert et al. 2008). Currently, nine species of Microstomidae have been reported exclusively from freshwaters while one (*M. lineare*) inhabits both limnic and brackish waters to 7‰. Further broad sampling efforts in different parts of the world will be required to illuminate the origin and distribution patterns of freshwater Microstomidae.

### Morphological Taxonomy

Morphological identification of flatworms in general is often dependent on the sexual anatomy, and species identification of Macrostomorpha specifically is heavily or entirely based on the morphology of the reproductive organs (e.g. Nasonov 1935; Ax & Armonies 1987; Faubel & Cameron 2001). Some authors have argued that species of Microstomidae with unknown sexual anatomy are incompletely described and should not be considered valid (Faubel 1984). Unfortunately, specimens of Microstomidae are most often immature when collected due to very short seasons of sexual reproduction, and thus scientists are often reluctant to describe new species (e.g. Yanoviak 2001; Delp 2002; Janssen et al. 2015). Ignoring or invalidating immature Microstomidae leads to large parts of biodiversity left unexplored in a group of animals that are ubiquitous, abundant (e.g. Martens & Schockaert 1981, Reise 1984) and ecologically important (Zeppilli et al. 2015) and prevents the discovered taxa from being included in future research or conservation efforts (Jörger & Schrödl 2013). Further, invalidating asexually reproducing immature species assumes that all species must have sexual anatomy, an assumption that is not necessarily legitimate. Instances of obligate asexuality are widespread throughout the animal kingdom (reviewed in Schön et al. 2009), and animal taxa already possessing the ability to reproduce asexually can give rise to asexual lineages without sexual organs, as has been demonstrated in e.g. aphids (Simon et al. 2002), ostracods (Butlin et al. 1998) as well as numerous other meio- and macrofaunal taxa (Simon et al. 2003).

In order to facilitate future research, we created a key to Microstomidae that is not dependent on the reproductive system so that species may be distinguished even when immature. As the taxonomic literature is often old and may be difficult to acquire, we further summarize the morphological and molecular taxonomy of all currently accepted species within Microstomidae. Important distinguishing morphological characters include: shape of anterior end (blunt, conical, proboscis, rounded; Fig. 11); shape of posterior end (blunt, pointed, rounded, spatulate, tail-like; Fig. 12); size and shape of ciliary pits (rounded, bottle-shaped; Fig. 13); length of preoral intestine (defined as long if extending anterior to the brain, medium if reaching to level of brain, and short if terminating posterior to level of brain; Fig. 14); body pigmentation; presence, location and color of pigmented eyespots; presence and location of rhabdites; presence, pattern and location of adhesive papillae; as well as other characters specific to individual species (e.g. the lateral intestinal bulges of *M. mundum*; the biparte pharynx of *M. breviceps*; Fig. 15). Table S3 summarizes the morphological characters for each species.

**Fig. 11:**
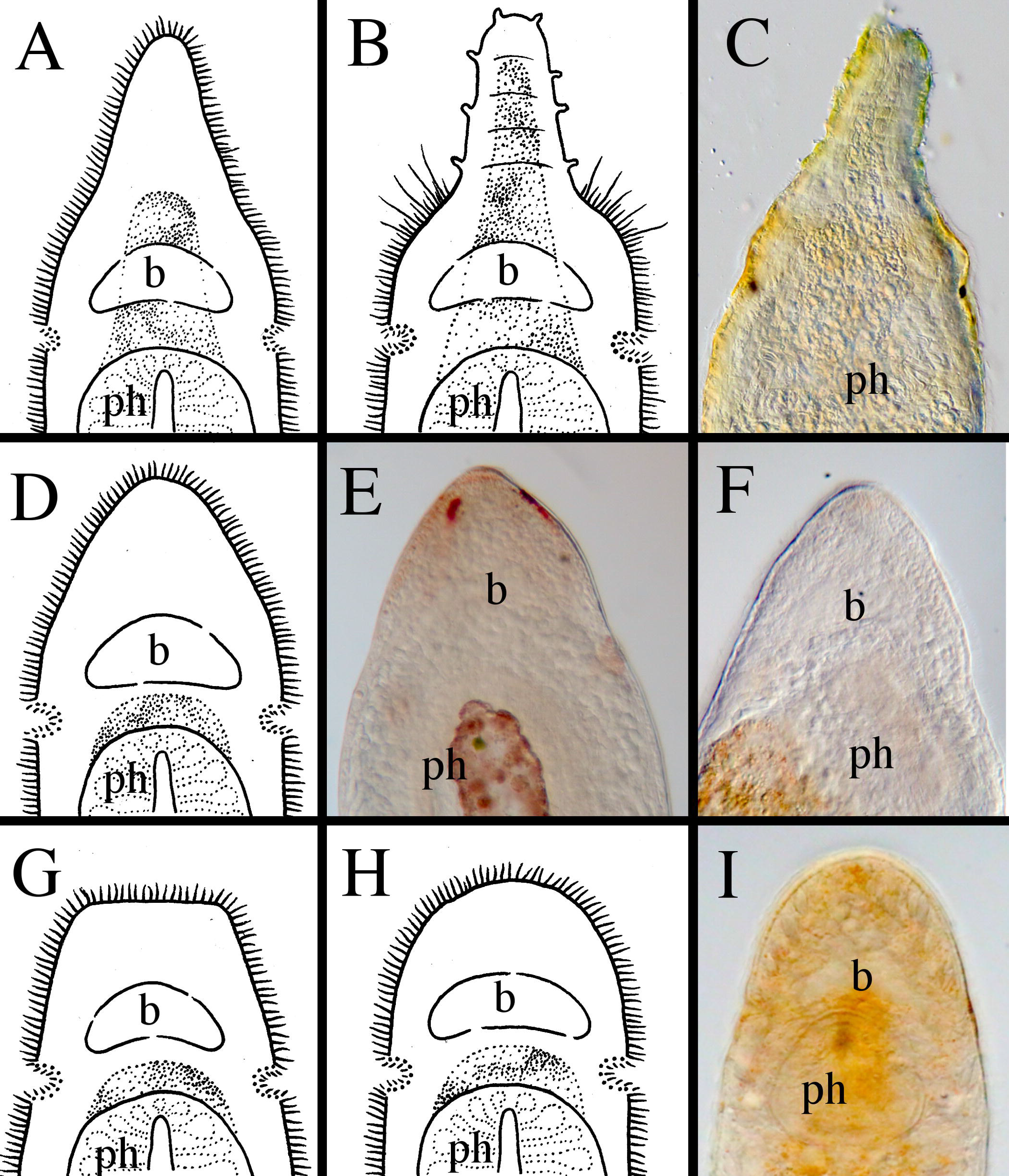
Different shapes of the anterior ends within *Microstomum*. A. Drawing depicting the elongated frontal end of *M. trichotum*. B. Drawing depicting an anterior proboscis. C. Micropictograph of the proboscis of *M. compositum*. D. Drawing depicting a conical anterior. E. Micropictograph of the conical anterior of *M. lineare*. F. Micropictograph of the conical anterior of *M. bispiralis*. G. Drawing depicting a blunt anterior. H. Drawing depicting a rounded anterior. I. Micropictograph depicting the rounded anterior of *M. septentrionale*. b brain; ph pharynx

**Fig. 12:**
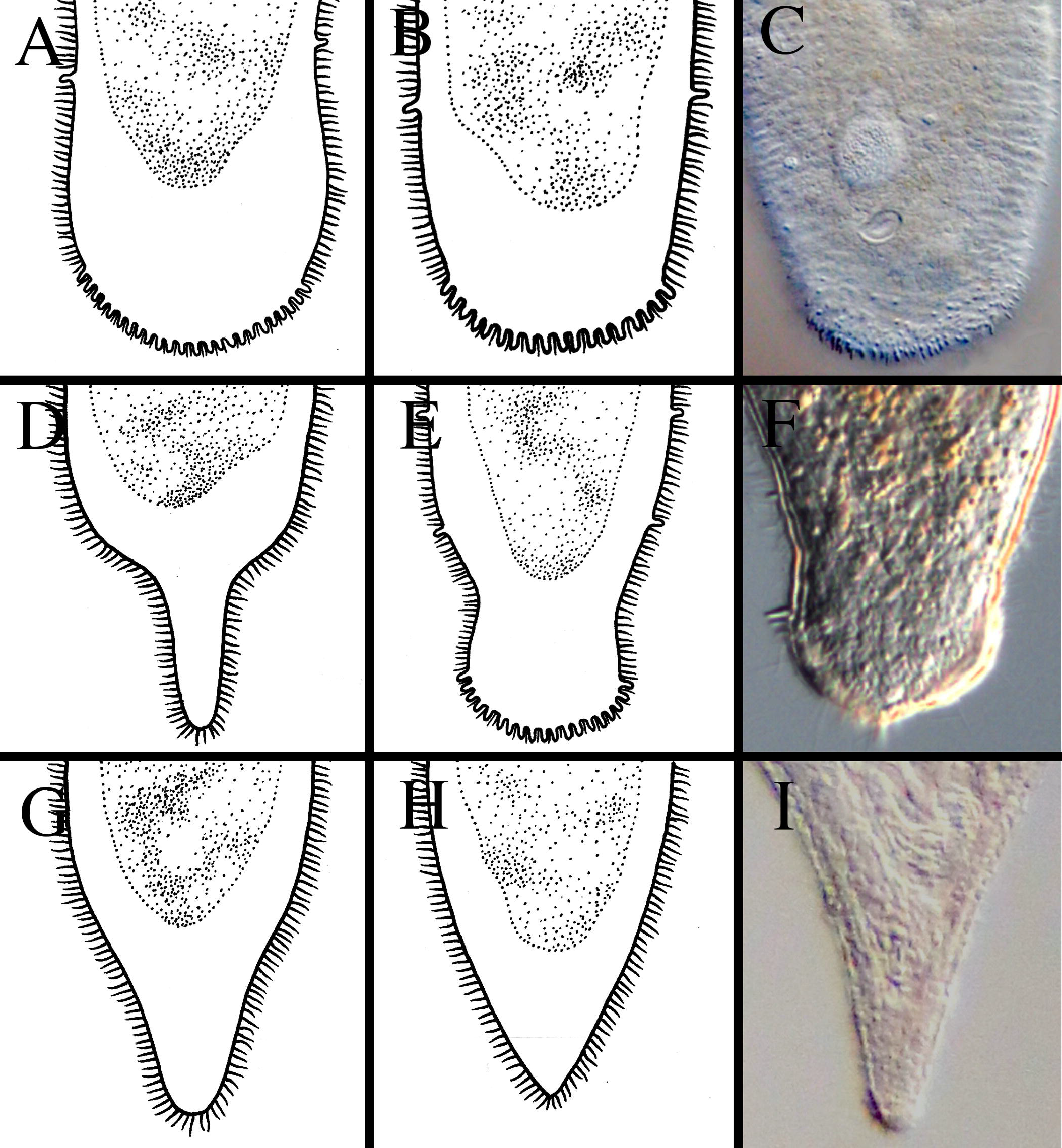
Different shapes of the posterior ends within *Microstomum*. A. Drawing depicting a rounded posterior. B. Drawing depicting a blunt posterior. C. Micropictograph of the blunt posterior of *M. edmondi*. D. Drawing depicting a thin-tail posterior of e.g. *M. giganteum*. E. Drawing depicting a spatulate posterior. F. Micropictograph of the spatulate posterior of *M. lotti*. G. Drawing depicting a tail-like posterior. H. Drawing depicting a pointed posterior. I. Micropictograph depicting the pointed posterior of *M. bispiralis*.

**Fig. 13:**
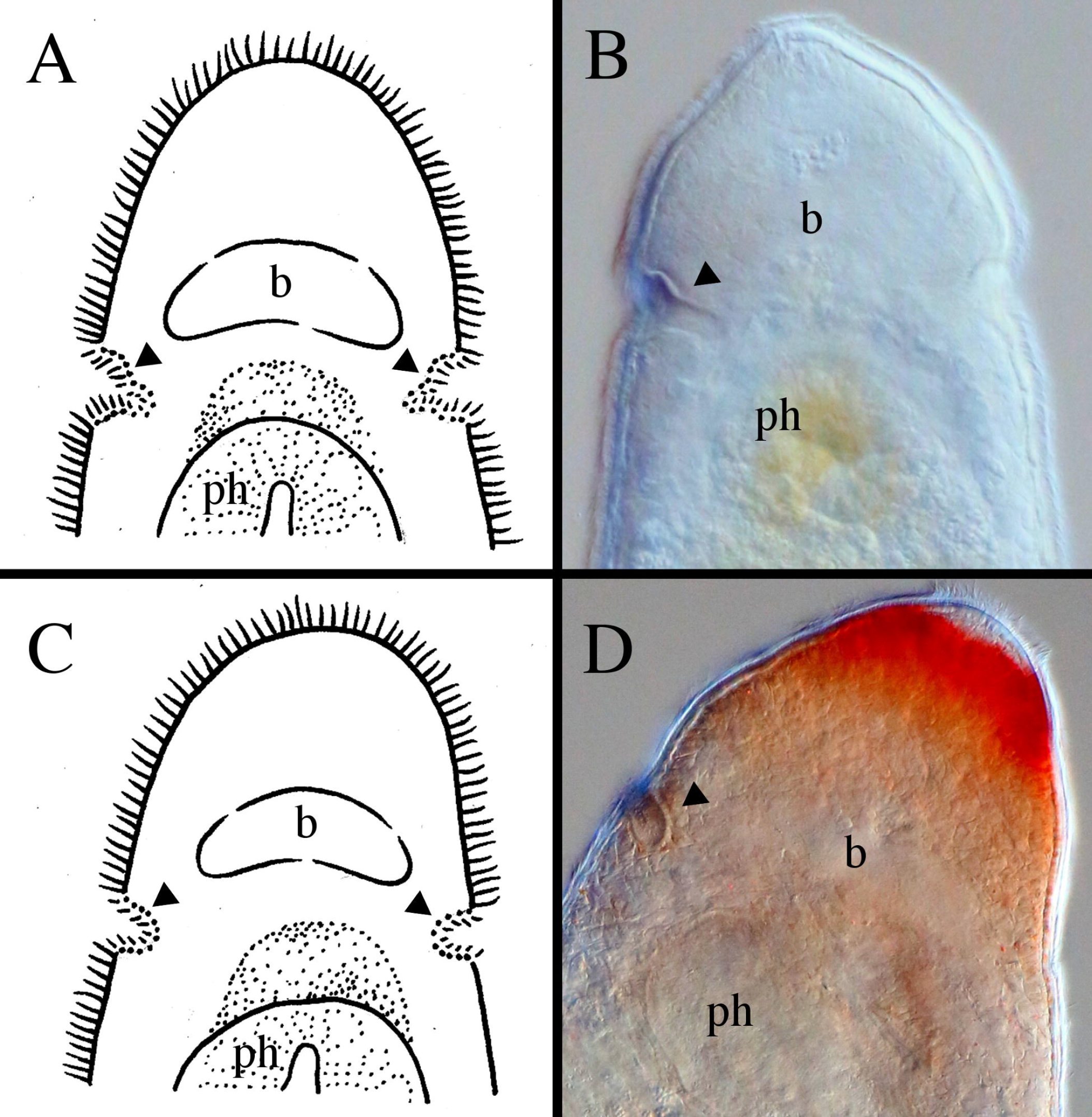
Different shapes of the ciliary pits within *Microstomum*. A. Drawing depicting bottle-shaped ciliary pits. B. Micropictograph of the bottle-shaped ciliary pits of *M. edmondi*. C. Drawing of rounded ciliary pits. D. Micropictograph of the rounded ciliary pits of *M. rubromaculatum*. Arrows indicate ciliary pits; b brain; ph pharynx

**Fig. 14:**
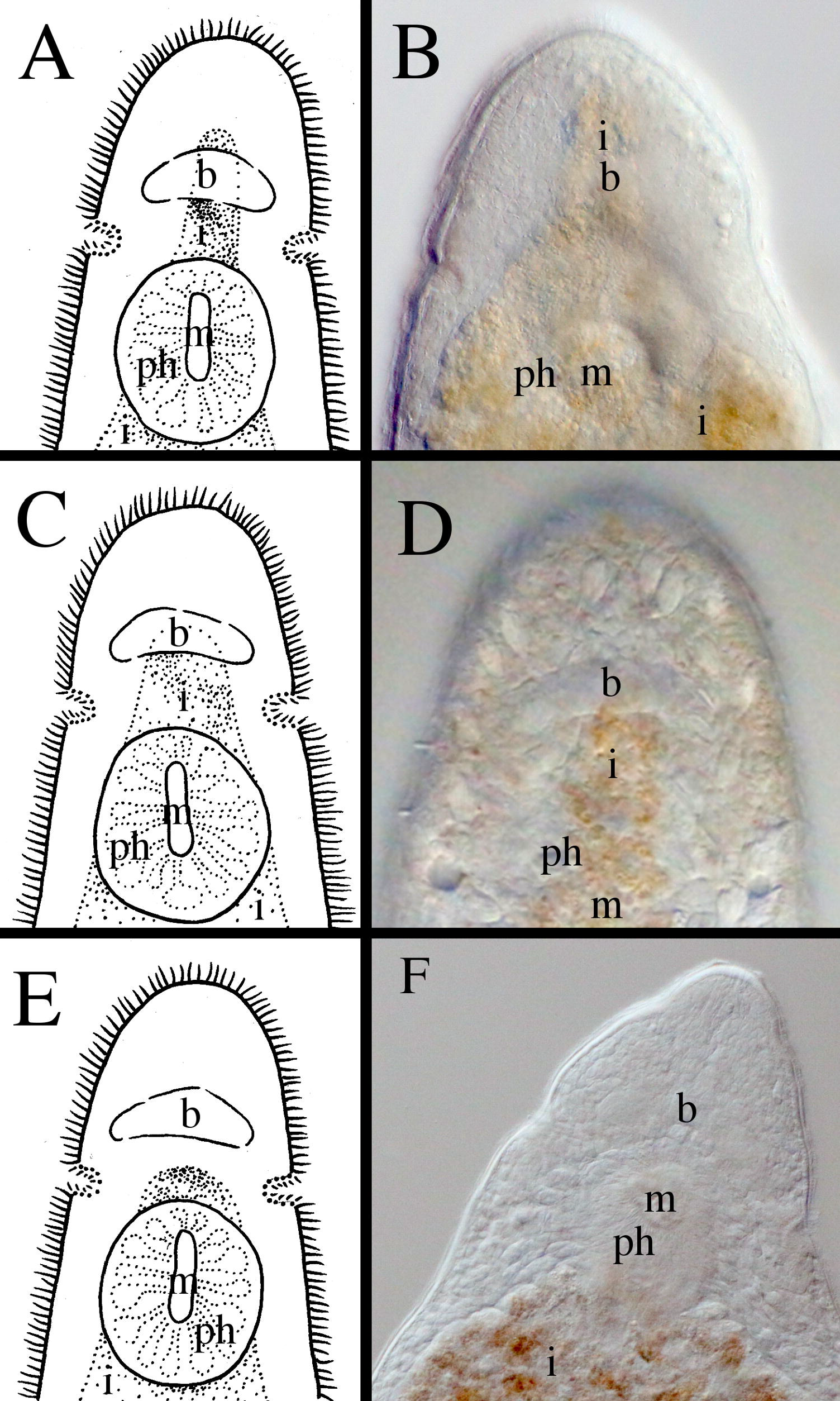
Different lengths of the preoral intestine within *Microstomum*. A. Drawing depicting a long preoral intestine that extends anterior to the brain. B. Micropictograph of the long preoral intestine of *M. inexcultus* sp. nov. C. Drawing depicting a medium preoral intestine that terminates at the level of the brain. D. Micropictograph of the medium preoral intestine of *M. septentrionale*. E. Drawing depicting a short preoral intestine that terminates posterior to the brain. D. Micropictograph of the short preoral intestine of *M. bispiralis*. b brain; i intestine; m mouth; ph pharynx

**Fig. 15:**
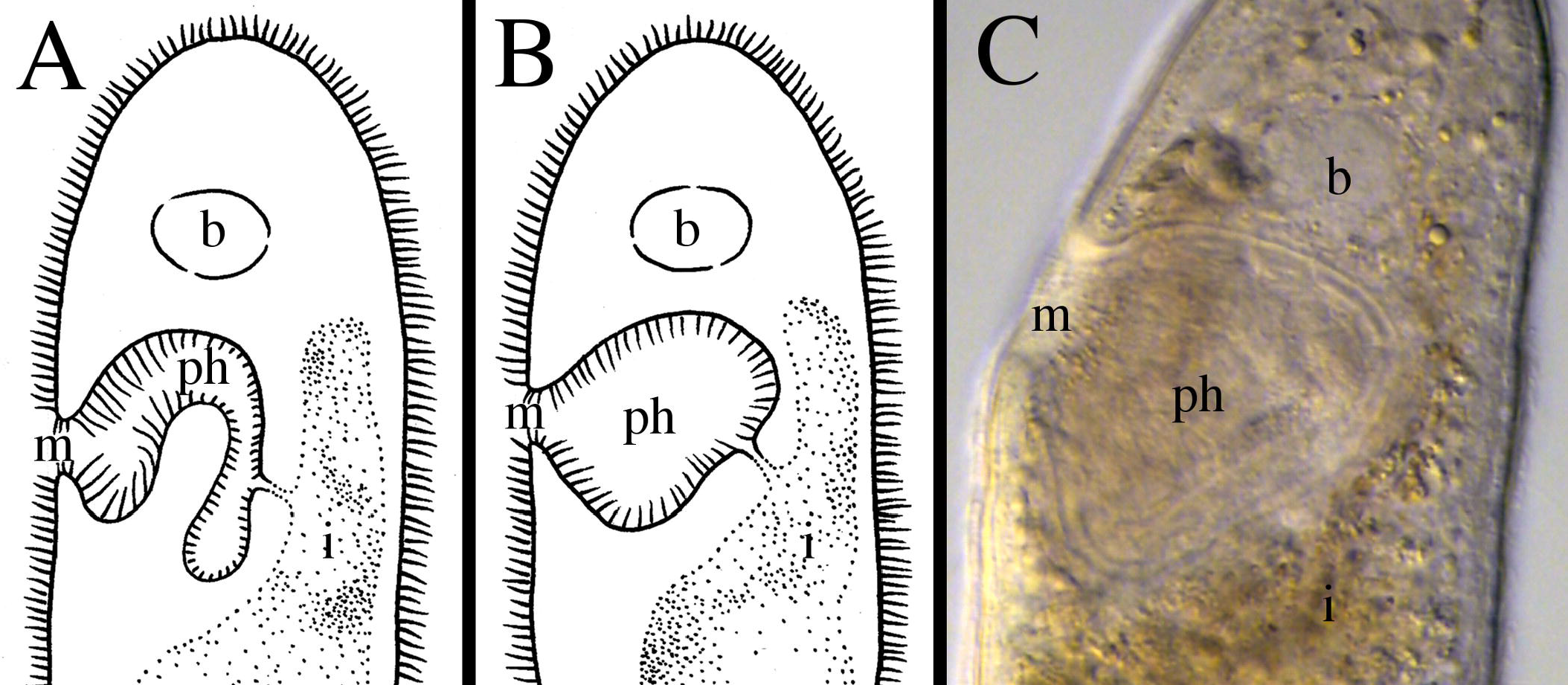
Different shapes of the pharynx within *Microstomum*. A. Drawing depicting the characteristic biparte pharynx of e.g. *M. breviceps*. B. Drawing depicting a typically-shaped pharynx of *Microstomum* C. Micropictograph of the typically-shaped pharynx of *M. bispiralis*. b brain; i intestine; m mouth; ph pharynx

## Taxonomic Summary

### Fam. Microstomidae LUTHER, 1907

Macrostomida with intestine extending anterior to the mouth (preoral intestine). Ciliated sensory pits present at frontal end, typically around level of brain.

Frontal glands present surrounding pharynx. Asexual reproduction through fissioning occurring. Marine, brackish and freshwater habitats; two genera.

Type: *Microstomum* SCHMIDT 1848

### Gen. *Microstomum* Schmidt 1848

Microstomidae with ciliary pits at frontal end and preoral intestine. Two excretion openings in the anterior body. Rhabdites and nematocysts present in some species. Asexual reproduction through fissioning as well as sexual reproduction present. Sexual anatomy including one or two testes beside or behind the intestine; one or two ovaries consisting of one or more lobes and containing a central oocyte and a peripheral layer of adipose cells. Body size of solitary individual 0.6-4mm, with chains comprising up to 18 zooids and 15 mm.

Type species: *Microstomum lineare* (Müller, 1773).

43 species: 33 marine, 9 limnic, and 1 inhabiting both brackish (up to 7‰) and freshwaters.

*Species Key:*

|  |  |  |
| --- | --- | --- |
| 1. | Pigmented Eyes Present | 2. |
|  | Pigmented Eyes Absent | 13. |
| 2. | Pigmented eyes red | 3. |
|  | Pigmented eyes dark, not red | 9. |
| 3. | Adhesive papillae present at the posterior end of the animal | 4. |
|  | Posterior adhesive papillae not present | 7. |
| 4. | Adhesive papillae present at the anterior end of the animal | 5. |
|  | Anterior adhesive papillae not present | 6. |
| 5. | Eyespots marginal dots close to anterior tip; body color reddish; preoral intestine to level of brain | *M. gabriellae* |
|  | Field of red pigmentation forming broad transverse band at anterior end; preoral intestine small, not reaching to brain | *M. rubromaculatum* |
| 6. | Paired red eyespots as small dots located at lateral margins of the body at level of brain, posterior to small ciliary pits | *M. laurae* |
|  | Single medial eyespot, oval shaped, located anterior to the brain; intestine with lateral lobules and bulges | *M. groenlandicum* |
| 7. | Eyespots present as red stripes at dorso-anterior end | *M. lineare* complex** |
|  | Eyespot not so | 8. |
| 8. | Body color yellow to orange; eyespots at level of ciliary pits; rhabdites present concentrated at body ends | *M. spiriferum* |
|  | Body yellow-green with two small eyespots at frontal margins; posterior end with stiff cilia twice as long as locomotory cilia; rhabdites present across body | *M. bioculatum* |
| 9. | Anterior end enlarged to form a proboscis with papillae and sensory cilia | 10. |
|  | Without anterior proboscis | 11. |
| 10. | Sensory cilia even spaced along lateral margins of body; eyespots posterior to mouth | *M. proliferum* |
|  | Without evenly spaced lateral sensory cilia; eyespots present just anterior to mouth | *M. compositum* |
| 11. | Preoral intestine short, not quite reaching level of brain; ciliary pits small; without adhesive papillae | *M. dermophthalmum* |
|  | Not so | 12. |
| 12. | Eyes with distinct crescent lens present at frontal end; posterior end tapering and rounded; marine species | *M. melanophthalmum* |
|  | Eyes with distinct crescent lens present anterior to ciliary pits; posterior end tail-like; freshwater species | *M. paradii* |
| 13. | Adhesive papillae present at the anterior end of the animal | 14. |
|  | Anterior adhesive papillae not present | 28. |
| 14. | Posterior end forming a spatulate tail plate set off by a constriction in the body width | 15. |
|  | Posterior end not so | 17. |
| 15. | Ciliary pits small and shallow; preoral intestine reaching anterior to brain; pharynx biparte with longer anterior cilia and shorter posterior | *M. breviceps* |
|  | Not so | 16. |
| 16. | Adhesive papillae on frontal end situated in a row ~30 μm from the anterior tip of the animal | *M. lotti* |
|  | Rhabdites heavily concentrated at anterior end above pharynx | *M. ulum* |
| 17. | Preoral intestine short, terminating posterior to level of brain | 18. |
|  | Preoral intestine extending to brain or anterior | 22. |
| 18. | Rhabdites present | 19. |
|  | Rhabdites absent | 21. |
| 19. | Anterior end enlarged to form a proboscis, non ciliated but with sensory bristles | *M. album* |
|  | Anterior end not so | 20. |
| 20. | Anterior end bluntly rounded; field of red pigmentation forming broad transverse band at anterior end | *M. rubromaculatum* |
|  | Body evenly light reddish-brown; anterior end rounded and set apart by slight constriction at ciliary pits | *M. schultei* |
| 21. | Body color brownish yellow with numerous anterior black dots; anterior end pointed and separated by distinct constriction at ciliary pits | *M. punctatum* |
|  | Body colorless; anterior end long and rounded; with numerous adhesive papillae on posterior end | *M. marisrubri* |
| 22. | Preoral intestine extending anterior to brain | 23. |
|  | Preoral intestine terminating at level of brain | 25. |
| 23. | Ciliary pits clearly discernable and bottle-shaped; adhesive papillae present in a row ventrally, ~25 µm from the anterior tip | *M. weberi* |
|  | Not so | 24. |
| 24. | Rhabdites very numerous at frontal end; adhesive papillae scattered along margins of entire body | *M. papillosum* |
|  | Rhabdites only sparsely present; few adhesive papillae present on anterior and posterior ends only | *M. afzelii* |
| 25. | Rhabdites present throughout body and extremely numerous at anterior end | 26. |
|  | Rhabdites not present or present at anterior ends only | 27. |
| 26. | Body colorless; field of adhesive papillae present at anterior tip of animal, longest at lateral sides and sloping upwards medially | *M. westbladi* |
|  | Body yellow with darker yellow to orange frontal end; adhesive papillae sparsely scattered over entire body | *M. septentrionale* |
| 27. | Adhesive papillae present at both body ends; pharynx biparte with longer anterior and shorter posterior cilia; paired stripes of rhabdite bundles on dorsolateral sides | *M. rhabdotum* |
| 28. | Rhabdites present | 29. |
|  | Rhabdites absent | 37. |
| 29. | Posterior end tail-like | 30. |
|  | Posterior end rounded or blunt, not tail-like | 31. |
| 30. | Rhammite glands colored red; ciliary pits deep and narrow | *M. jenseni* |
|  | Intestine with lateral bulges and smaller lobules | *M. mundum* |
| 31. | Ciliary pits large and very distinct, bottle shaped | 32. |
|  | Ciliary pits not so | 33. |
| 32. | With ochre frontal end; preoral intestine very short, hardly extending past mouth | *M. curinigalettii* |
|  | Body colorless; preoral gut extending anterior to brain | *M. edmondi* |
| 33. | Preoral intestine extending anterior to brain | 34. |
|  | Preoral intestine terminating at level of brain | 36. |
| 34. | Pharynx divided into two parts by annular fold with longer anterior and shorter posterior cilia | *M. davenporti* |
|  | Pharynx not so | 35. |
| 35. | Body red with yellow rhammite glands | *M. mortenseni* |
|  | Body colorless; rhabdites concentrated at anterior end | *M. spiculifer* |
| 36. | Anterior end long and tapered, resembling a proboscis; body white | *M. trichotum* |
|  | Body colorless and may have ± dark grey pigment spots in some individuals; anterior end bluntly rounded | *M. lucidum* |
| 37. | Preoral intestine extending anterior to brain | 38. |
|  | Preoral intestine terminating at or before level of brain | 40. |
| 38. | Adhesive papillae present at posterior tip and along up to ~1/5 of the posterior body | *M. inexcultus* |
|  | Adhesive papillae absent | 39. |
| 39. | Ciliary pits deep; anterior end conically pointed; posterior end tail-like with long sensory cilia | *M. canum* |
|  | Body ends blunt; bluish color | *M. caerulea* |
| 40. | Rounded posterior end with adhesive papillae | *M. crildensis* |
|  | Posterior adhesive papillae not present | 41. |
| 41. | Preoral intestine reaching to level of brain | *M. lineare* |
|  | Preoral intestine terminating before level of brain | 42. |
| 42. | Animal comparatively large, single zooid to 4 mm, 18.5 mm maximum for 8 zooids; posterior end rounded or with thin-tail | *M. giganteum* |
|  | Maximum body length 4 mm, 8 zooids; body transparent grey to light yellow with tapered pointed posterior end | *M. bispiralis* |

\*\**M. lineare* complex refers to the species that are distinguished from *M. lineare* primarily on the basis of DNA information, specifically *M. artoisi*, *M. tchaikovskyi*, *M. zicklerorum* (Atherton & Jondelius, 2018b)

*List of Currently Recognized* Microstomum *Species in Alphabetical Order*

#### 1. *Microstomum album* (ATTEM 1896)

##### Synonyms

*Alaurina alba* Attems 1896, pg 56

Type: Not recorded

Type Locality: North Sea, Helgoland Habitat: Marine, pelagic, phytal

Brinkmann 1905 – Denmark, Frederikshavn in Kattegat Graff 1913 – France, Gulf of Lion, Banyuls sur Mer

Body length of solitary animal to 1.6 mm, chains to 2.5 mm, 4 zooids. Body colorless or white, displaying yellow-orange intestine. Pigmented eyes absent. Anterior end smoothly tapering to conically pointed proboscis, non-ciliated but with tactile bristles and a ring of papillae. Posterior end rounded. Small ciliary pits present slightly anterior to mouth. Rhabdites present, numerous over entire body. Preoral intestine short. Adhesive papillae sparsely present along entire mid to posterior body and highly concentrated at posterior end.

Reproductive system with single testis and ovary. Stylet arched projecting into antrum masculinum. True prostatic vesicle present in addition to seminal vesicle present. Female system with ovary and short duct. Gonopores separate.

#### 2. *Microstomum afzelii* sp. nov.; Fig. 3

Type: Asexual, MTP LS 700; GenBank accession 18S KP730494

Type Locality: Sweden, Saltö, 58°52’29’’N 11°08’41’’ E, 10 cm

Habitat: Marine, eulittoral sand

##### GenBank Accession

18S – KP730494

Body length 0.8 mm, 2 zooids. Body colorless, without pigmented eyespots. Body shape elliptical with rounded anterior and posterior ends. Ciliary pits weakly present, small and difficult to discern. Adhesive papillae and rhabdites sparsely present. Nematocysts particularly concentrated at posterior end. Preoral intestine long, extending anterior to brain.

Reproductive system unknown.

#### 3. *Microstomum artoisi* Atherton & Jondelius 2018; Fig. 17D

Type: Asexual, SMNH-Type-9022; GenBank accession 18S MH221276, CO1 MH221384, ITS MH221478

Type locality: Belgium, Hoge Kemper Park, 51°00’03’’N 5°40’44’’E, 10 cm depth Habitat: Freshwater phytal, 5-10 cm depth

##### GenBank Accession

18S – MH221275 – MH221276
CO1 – MH221383 – MH221384
ITS – MH221477 – MH221478

##### Molecular Diagnosis

COI gene region with reference to GenBank Accession MH221288: 39-A; 42-G; 57-A; 81-C; 114-A; 180-T; 195-C; 207-C; 216-G; 234-C; 240-G; 321-G; 390-C; 409-A; 612-T

ITS1, 5.8S, ITS-2 gene region with reference to GenBank Accession MH221390: 56-A; 383-T; 839-A; 961-T

##### Morphological Description

Body length to approximately 3.5 mm, maximum 6 zooids. Body color greyish or slightly tinged pinkish clear. Anterior end abruptly conical, posterior end tail-like without adhesive papillae. Sensory bristles present and ciliary pits present.

Two red eyestripes present at dorso-anterior end. Amount of eyespot pigmentation in some individuals very bright, almost merging medially to form a dorsal. Preoral intestine short.

Reproductive system not seen.

Notes: See notes of *M. lineare*.

#### 4. *Microstomum bioculatum* Faubel, 1984

Type: Not deposited

Type locality: North Sea, Sylt Island, off Keitum

Habitat: Marine, eulittoral and superlittoral lentic upper mudflats and salt meadows.

Body length to 1.4 mm, maximum 4 zooids. Body yellow-green with two very small red eyespots at frontal margins. Ends rounded; posterior with long, stiff cilia that are twice the length of the locomotory cilia. Rhabdites present. Ciliated pits, preoral intestine and adhesive papillae all apparently absent.

Reproductive system not seen.

Notes: The absence of the preoral intestine and, especially, ciliated pits is extremely unusual, as both are defining characters of the genus. Additionally, there seems to be some confusion regarding the preoral intestine, since the original written description by Faubel (1984) specifically precludes its presence (“Ein Kopfdarm und typische Einschnürungen des Körpers in Höhe des Gehirns fehlen”), but Faubel’s original drawing of the species distinctly includes a preoral intestine that extends anterior to the brain.

#### 5. *Microstomum bispiralis* Stirewalt, 1937; Figs. 16, 17E

Type: Deposition not recorded

Type locality: near University of Virginia, Virginia, USA Habitat: Freshwater pools

This paper: Sweden, Finland

GenBank Accession:

18S – XXX
CO1 – XXX

Maximum 8 zooids, 4 most common, with body length to 4 mm. Following laboratory cultures, individuals can be transparent-grey to bright yellow. Pigmented eyespots absent. Anterior end short conically pointed. Posterior end tapered to a point. Ciliary pits present anterior to mouth, each shaped as a tube extending posteriorly at 45° angle and ending in an enlarged spherical pocket. Pharynx ciliated, and occupies ⅓ to ½ length of anterior zooid. Preoral intestine short. Adhesive papillae and rhabdites absent. Nematocysts may be present in some specimens.

Nervous system and cerebral ganglion as typical for the genus; consisting of two anterior ganglia connected by a short and wide transverse commissure. A dorsolateral and a ventrolateral nerve extend posteriorly from each ganglion.

Ventrolateral nerves are connected by a commissure ventral to the pharynx. One nerve extends laterally from each ganglion to the ciliary pit and another from each ganglion to branch profusely in the anterior end of the body.

Protanderous hermaphrodites, with male formation occurring in laboratory cultures during January and February. Male genital system consisting of two compact and spherical testes, one developing earlier than the other. Vasa deferentia extending posteriorly from each testis to empty dorsolaterally into the vesicula seminalis. Vesicula seminalis shaped as a small sphere, located ventral and posterior to the intestine; glandular and surrounded by a heavy muscular sheath of the copulatory organ. Copulatory stylet a smooth spiral sharply bent in two planes, 120 μm long, widened at the base and drawn out distally to a sharp point.

Female reproductive system includes a single ovary lying slightly anterior to the middle of the zooid, ventral to the intestine. Numerous single-celled glands open to the vagina, which opens to a female pore at the beginning of the posterior ⅓ of the zooid.

Somatic chromosome number 16 (n=8)

Notes: This is the first time that *M. bispiralis* is reported from Sweden or anywhere else outside of the original location. The specimens collected in Finland and Sweden matched the original descriptions, including the male stylet and other sexual anatomy that was found in three individuals (Fig. 16C). The very great distance between the locations does suggest the possibility of cryptic species or much larger dispersal abilities than has previously been shown in any species of *Microstomum* (see Atherton & Jondelius 2018b), although true support for either scenario must be deferred until DNA sequence data can be attained from specimens collected at the type locality.

**Fig. 16.**
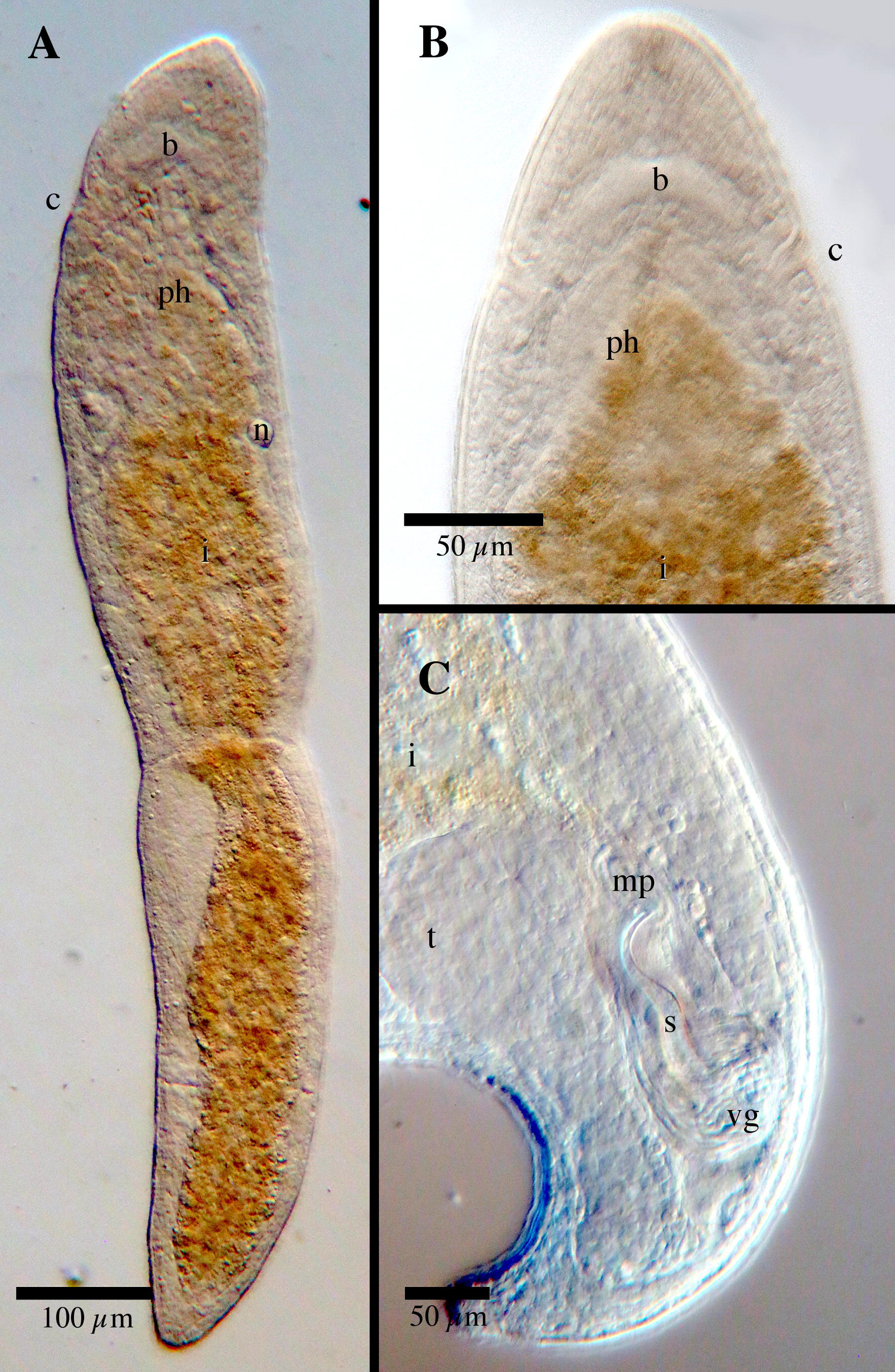
*Microstomum bispiralis*; specimen from Sweden. A. Micropictograph of whole body B. Anterior end. C. Posterior end with focus on sexual anatomy. b brain; c ciliary pits; i intestine; mp male pore; n nematocyst; ph pharynx; s male stylet; t testis; vg vesicula granulorum

The absence of adhesive papillae and rhabdites combined with a short preoral intestine is a unique set of characters that separates this species from nearly all of the other currently accepted *Microstomum* species. *M. bispiralis* is most similar to *M. lineare* and *M. lineare*-like or cryptic species (including *M. artoisi, M. giganteum*, *M. tchaikovskyi*, and *M. zicklerorum*), which in part is due to the very wide morphological variability reported for *M. lineare*. However, *M. bispiralis* is typically smaller with a slightly higher body width:length ratio, has a shorter preoral intestine, and always and entirely lack the pigmented eyestripes. Further, all species within the *M. lineare* complex are separated through 18S and CO1 gene evidence (see Figs. 1, 2), while the shape and size of the male stylet is different between *M. bispiralis*, *M. giganteum* and *M. lineare*.

#### 6. *Microstomum breviceps* Marcus, 1951

Lectotype: Asexual, sectioned, SMNH-Type-XXXX; E. Marcus leg.** Type locality: Brazil, São Sebastião

Habitat: Marine, between algae, mainly *Sargassum stenophyllum*

Maximum 3 zooids, 0.7 mm. Color and pigmented eyes unknown as worms were only described based on preserved state. Anterior end rounded, posterior spatulate. Adhesive papillae present anteroventrally above level of the brain, at the posterior tip and in the zone of division in mature zooids. Ciliated pits small. Rhabdites occur throughout the body. Preoral intestine extends anterior to brain.

Mouth circular. Pharynx divided into two parts by an annular fold, with longer cilia in the anterior part and shorter cilia in the posterior. Glands around the pharynx may be pale pink in color. Intestine unciliated.

Reproductive organs not seen.

Notes: *Microstomum breviceps* is similar to *M. davenporti* in that both have similar patterns of adhesive papillae, a long preoral intestine and a characteristic divided pharynx with long and short cilia and with an annular fold. *M. breviceps* can be separated from *M. davenporti* by its body shape: the anterior end of *M. breviceps* is truly round instead of tapering blunt, and the posterior spatulate tail-plate is very distinct from the rest of the body. Other differences can be found in the high concentration of rhabdites surrounding the oval mouth in *M. davenporti*, and the very large distances separating their locations (São Sebastião, Brazil and Woods Hole Massachusetts, USA for *M. breviceps* and *M. davenporti*, respectively).

**A type specimen was not previously designated for *M. breviceps.* However, two original specimens were deposited at the Swedish Museum of Natural History, and thus, following the rules of the International Code of Zoological Nomenclature, we hereby designate one sectioned asexual specimen (formerly SMNH108136, now SMNH-Type-XXXX) as lectotype and one whole mount asexual specimen (SMNH108137) as paralectotype for *Microstomum breviceps* MARCUS, 1951.

#### 7. *Microstomum caerulescens* (Schmarda, 1859)

Synonyms:

*Strongylostomum caerulescens* - Schmarda 1859
*Strongylostomum coerulescens* - Parádi 1881, pg 311; Diesing 1862, pg 235; Graff 1882, pg 253
*Typhlomicrostomum coerulescens* - Diesing 1862
*Microstomum coerulescens* - Graff 1882, pg 253

Type: Not recorded

Type locality: Jamaica, Freshwater lakes near Kingston

Habitat: Freshwater lakes

Body small, 0.67 mm; two zooids. Anterior and posterior ends blunt, almost flat. Bluish color; eyespots not recorded. Preoral intestine extends to close to the anterior end. Adhesive papillae, rhabdites and ciliary pits not recorded.

Reproductive system not observed.

Notes: Ciliary pits were not described for this species, nor were they included in Schmarda’s (1859) drawing. However, the original description occurred at a very early time period, and positively identifying ciliary pits may be difficult even with modern equipment when they are small and/or shallow. The importance of the character for this genus (as first argued by Uljanin 1870; Graff, 1882) suggests that the presence or absence of ciliary pits for this species should remain in question until they can be re-examined. Even without absolute knowledge of the adhesive papillae, ciliary pits, and reproductive system, the combination of blue body color, absence of pigmented eyespots, large pre-oral intestine, blunt body ends and freshwater habitat is enough to separate this species from all other currently described species of *Microstomum*.

#### 8. *Microstomum canum* (Fuhrmann 1894)

Synonyms:

*Microstoma canum* Fuhrmann 1894, pg 232

Type: Not recorded

Type locality: Switzerland, Basel, Augustinerholzbach

Habitat: Freshwater

Graff 1913 – France

Body length to 2.0 mm, 8 zooids. Body grey. Eyes absent. Anterior end long and tapering to a point. Posterior end pointed, with long cilia but without adhesive papillae. Ciliary pits deep, present at level of mouth, surrounded by long cilia. Preoral intestine extends anterior to brain. Rhabdites and nematocysts not seen.

Reproductive system unknown.

Notes: Hofsten (1912) and Meixner (1915) postulated that *M. canum* is a synonym of *M. lineare* based on variability of the latter. While the wide morphological variations of *M. lineare* undoubtedly creates difficulty in absolutely ruling out synonymy, potential differences between the two species exist in the extension of the preoral intestine anterior to the brain, a more posterior position of the ciliary pits and, perhaps, a more pointed anterior end (based Furhmann’s original drawing) for *M. canum*.

#### 9. *Microstomum compositum* (Metschnikov 1865); Fig. 18F

Synonyms:

*Alaurina composita* Metschnikov 1865, pg 178 *Alauretta viridirostrum* Mereschkowsky 1879, pg 35 *Alaurina viridirostrum* Graff 1882, pg 261 *Alaurina claparedii* Graff 1882, pg 262 *Dinophilis simplex* Verrill 1892, pg 485
Type: deposition not recorded Type Locality: Germany, Helgoland, Coast of Skye-Hebrides Habitat: Marine pelagic or planktonic
Brinkmann 1905 – Denmark, Skagens Rev, Horns Rev Gamble 1893 – United Kingdom, Island of Skye Graff 1913 – United States, Rhode Isalnd, Newport Hofker 1930 – Netherlands, Den Helder, IJsselmeer This paper: Sweden, Fiskebäckskil, Kristineberg Sven Lovén Center for Marine Research, 58°14′59′′ N, 11°26′45′′

GenBank Sequences

18S – XXX CO1 – XXX

Body length of solitary animal to 0.9 mm, chains to 3.0 mm. Maximum 10 zooids, 4 most common. Body color light yellow with greenish anterior (proboscis). Black pigmented eyespots present. Anterior end sharply decreasing in width to form distinct proboscis, with sensory cilia, papillae and numerous circular folds, and denoted by paired transverse tufts of cilia at base. Posterior tip with thin tail and terminal sensory cilia. Preoral intestine extending anterior to brain. Intestine non-ciliated, often with diatoms and protozoa. Rhabdites may be present in the anterior end.

Male reproductive system including a single testis, which may be bilobed in some specimens. Stylet weakly arched connecting to seminal vesicle.

Female system unknown.

Notes: Brinkmann (1905, pg 66) argued that both *Alaurina viridirostrum* and *A. claparedii* were identical to this species, stating that the very few differences were due to errors derived from outdated methods of investigation and/or a lack of sufficient specimens.

#### 10. *Microstomum crildensis* Faubel, 1984; Fig. 18E

Type: Deposition not recorded

Type locality: Germany, Jade Busen, Crildumer Siel Habitat: Marine, eulittoral

This paper: Sweden, Fiskebäckskil, Kristineberg Sven Lovén Center for Marine Research, 58°14′59′′ N, 11°26′45′′; Stömstad, Saltö, 58°52’29’’N 11°08’41’’ E

GenBank Accession:

18S – XXX CO1 – XXX

Body to 1.5 mm, maximum 4 zooids. Solitary animal 0.9 mm. Body colorless; pigmented eyes absent. Ends rounded, posterior with adhesive papillae. Constriction at level of brain at ciliary pits. Preoral intestine short, not quite reaching level of brain. Rhabdites and nematocysts absent.

Single testis, located at the very caudal end of the animal. Vesicula granulorum connected to a 32 μm long stylet, shaped as a curved tube of approximately consistent width and with distal terminal opening. Single ovary present.

Notes: Three specimens identified as *Microstomum crildensis* were collected from Swedish Skagerrak coast. Unfortunately, none of these specimens displayed sexual organs, and so the reproductive system—most importantly the male stylet—could not be used to corroborate identification. However, in every other respect, these specimens matched the original description and drawings, including: overall body shape including a body constriction at the ciliary pits, presence adhesive tubes on the posterior end only, absence of rhabdites, and small preoral intestine. The habitat— eulittoral muddy sediments of the North Sea—was also consistent.

Currently, the only other species of *Microstomum* with posterior adhesive papillae that entirely lacks rhabdites are *M. inexcultus* sp. nov., *M. marisrubri* sp. nov. and *M. punctatum.* M. crildensis can be distinguished from each of these species through:

*M. inexcultus* sp. nov. – long preoral intestine; constriction of the anterior body width near the ciliary pits; see also Figs. 1, 2
*M. marisrubri.* sp. nov. – shape and size of the male stylet; presence of anterior adhesive papillae and more numerous posterior adhesive papillae; see also Figs. 1, 2
*M. punctatum* – brown-yellow body coloring with anterior black spots; tail-like posterior end; limnic habitat

#### 11. *Microstomum curinigalletti* sp. nov.; Fig. 4

Type: Asexual specimen, SMNH-XXXX; GenBank Accession XXX

Type locality: Sweden, Fiskebäckskil, 58°14′59′′ N, 11°26′45′′ E, 20 cm depth Habitat: Marine, sublittoral phytal on algae

GenBank Accession:

18S – XXX CO1 – XXX

Body length 1.4 mm, 2 zooids. Anterior body end ochre to dark yellow-orange. Body otherwise colorless without pigmented eyespots. Anterior end conically rounded; posterior end blunt with numerous adhesive papillae. Ciliary pits large and bottle-shaped, located anterior to pharynx. Preoral intestine very short. Few rhabdites scattered across the body. Nematocysts present.

Reproductive system unknown.

#### 12. *Microstomum davenporti* Graff, 1911

Type: Deposition not recorded

Type locality: Woods Hole, Massachusetts, USA and Stamford, Connecticut, USA Habitat: Marine, on *Ulva* sp. and *Sargaassum* sp.

Body length to 1.5 mm; maximum 4 zooids. Body colorless or whiteish with a yellow-ocher intestine. Eyes absent. Anterior conically pointed; posterior end broad with numerous adhesive papillae. Ciliary pits present but very shallow. Rhabdites present, 12 μm, highly concentrated at the anterior end and around the oval mouth. Preoral intestine long. Pharynx divided into two parts by an annular fold, with longer cilia in the anterior part and shorter cilia in the posterior

Reproductive system not seen.

#### 13. *Microstomum dermophthalmum* Riedel, 1932

Type: Deposition not recorded.

Type locality: Engelskmandshavn, Godhavn, Greenland Habitat: Marine, coarse sand and sandy mud, 1-15 m depth

Steinböck 1938—West Greenland, sand and mud from 5-10 m depth; Iceland, low tide zone, coarse sand and algae, outer harbor of Ísafjör*∂*ur.

Riedl 1956 – Adriatic sea

Small, body length to 0.32 mm in solitary animal. Anterior end short conical; posterior end tapering to a rounded tip. Body colorless. Marginal eyes present very near the anterior end. Ciliary pits small and indistinct, at level posterior to marginal eyes. Rhabdites concentrated at anterior and posterior ends of the body. Adhesive papillae not recorded. Preoral intestine not quite reaching level of brain. Pharynx with 5.4 μm long cilia. Intestine unciliated.

Paired testes, sausage-shaped, obliquely lateral and posterior to ovary. Vesicula granulorum 20 μm long, 13 μm wide, strongly muscled. Male stylet strait tube with distal toothed thickening; small ~10-12 μm long (based on drawing).

Antrum masculine small, spherical. Genital pores separate.

#### 14. *Microstomum edmondi* Atherton & Jondelius 2018; Fig. 18A

Type: ♂, SMNH-Type-8890; GenBank accession MF185704

Type Location: Sweden, Fiskebäckskil, 58°14’52’’N, 11°27’05’’E, 25 cm Habitat: Marine benthic, 15-25 cm depth

GenBank Accession:

18S – XXX-XXX CO1 – MF185700 – MF185711

Body length to 1.7 mm, maximum 4 zooids. Conical pointed anterior end; blunt, almost flat posterior end with numerous 5-8 μm long adhesive papillae along rim. Ciliary pits large, bottle shaped. Body clear. Pigmented eyes absent. Dense field of cilia clearly covering epidermis. Preoral intestine extending anterior to brain. Mouth distinctly encircled by glands. Large rhabdite bundles to 45 μm long, primarily in the posterior end.

Protandrous hermaphrodite. Male reproductive system with single large testis. Vesicula seminalis circular to elliptical, 98 μm long, and containing the ends of numerous prostate glands in the distal part. Stylet approximately 67 μm long; shaped as an elongate, narrow funnel, slightly curved in one plane with a short, arched tip.

Female reproductive system with single ovary and gonopore. Eggs develop caudally.

#### 15. *Microstomum gabriellae* Marcus, 1950

Lectotype: Asexual, whole mount, SMNH-Type-XXX; E. Marcus leg.** Type location: Brazil, São Sebastião

Habitat: Marine, between algae, mainly *Sargassum stenophyllum*

Body length to 3 mm; maximum 7 zooids. Color reddish, with anterior marginal paired red eyespots. Anterior end short and conically rounded; posterior end rounded. Ciliated pits are thin and deep, located at level of posterior brain. Adhesive papillae present at the posterior of each well-developed zooid and more sparsely along the body. Rhabdites present, scarce and throughout the body. Nematocysts not seen. Pharyngeal nerve ring present, connected to brain by two short nerves. Preoral intestine reaching to level of brain.

Reproductive system based on original drawings (Marcus, 1950). Male system with paired round testes connecting separately to vesicular seminalis by vas deferens.

Stylet approximately 45-50 μm long, shaped as a tube with near consistent width throughout, broadly curved 180°.

**A type specimen was not previously designated for *M. gabriellae.* However, nine original specimens were deposited at the Swedish Museum of Natural History, and thus, following the rules of the International Code of Zoological Nomenclature, we hereby designate one whole-mount asexual specimen (formerly SMNH108140, now SMNH-Type-XXXX) as lectotype. The other eight originally deposited specimens (SMNH108138-9, SMNH108141-6) are thus designated as paralectotypes.

#### 16. *Microstomum giganteum* Hallez 1878

Type: Not recorded

Type Location: France, Lille (?)

Habitat: Freshwater in Europe, far less common than *M. lineare*

Hallez 1878, 1879 – France, Lille, Wimereux

Keller 1894 – Switzerland

Zabusov 1894 – Russia, Kazan

Zykov 1897 – Russia, in the viscidity of Moscow

Volz 1898 – Switzerland

Dorner 1902 – Germany, Königsberg

Markov 1904 – Russia, Kharkiv region, Udy River

Graff 1909 – Germany

1913 – Scotland, Denmark

Nasanov 1919 – Kirov oblast

Southern 1936 – Ireland, Clare, shore of Lough Atorick

Kostenko 1988 – Ukraine, Kiev, Okrung Reservoirs

Rogozin 2015 – Lipovka River falling into Lake Ilmen, 55°00′42′′ N, 60°09′20″ E; Lake Bol’shoe Miassovo, 55°10′39′′ N, 60°17′31″ E

Body large, single zooid to 4 mm, max length 18.5 mm for 8 zooids. Body whitish, grey or colorless with brown-red intestine. Pigmented eyespots absent. Anterior end rounded; posterior ends rounded with a thin tail. Adhesive papillae absent. Preoral intestine extending to anterior pharynx or very slightly past. Nematocysts present.

Male copulatory organ with an oblong seminal vesicle, 150-180 μm long. Stylet with three parts: 85 μm distal part inserted in a 30 μm mid part which is inserted in turn into a 40 μm long proximal part. Total length of the stylet to 180 μm, shaped as a spiral with 2.5 full turns. Caudal ovary with three follicles with triangular ovicels and 4-5 abortive eggs.

Notes: There was much discussion in the literature about whether *Microstomum giganetum* should be considered a “safe species” or a larger variant of *M. lineare*. The species name was reduced to a synonym by Luther (1960), who considered it to represent a pure line of *M. lineare* that had developed over a long period of asexual reproduction. *M. giganteum* eventually regained its independence after Kostenko (1988) carried out a comparative study of both forms and found several morphological differences, most importantly in the different structures of the reproductive system and male stylet, without intermediate or transitional forms.

*M. giganteum* differs from *M. lineare* through its larger size (8 zooid body length of ≥8.5 mm compared with 4-5 mm, respectively); lack of elongated tail, adhesive papillae and pigmented eyespots; pharyngeal glands restricted to oral cavity; and reproductive system including larger male stylet with three parts forming greater number of spiral turns (2.5 versus 1.5, respectively).

#### 17. *Microstomum groenlandicum* (Levinsen, 1879)

Synonyms:

*Microstoma groenlandicum* Levinsen 1879, pg 449
Type: Deposition not recorded Type Locality: Davis Strain, Disko Island, Greenland Habitat: Marine, among *Ulva* sp.
Gamble 1893 – United Kingdom, English Channel Graff 1882 – Greenland, Baffins Bay Graff 1905, 1913 – Norway, Bergen Meixner 1938 – United Kingdom, English Chanel Southern 1912 – Ireland, Clare Island Southern 1936 – Ireland, Blacksod Bay, New Harbor

Body length to 2 mm, with maximum 8 zoods. Body colorless, yellow or reddish. A single medial red eyespot anterior to the brain, oval shaped. Anterior tip of the body conically pointed. Posterior end long, tail-like with many adhesive papillae.

Rhabdites present throughout the body, concentrated at the anterior end. Small ciliary pits located at level between the brain and the mouth. Preoral intestine extending to the brain. Intestine with lateral lobules and bulges.

Testes not seen. Male stylet narrow, cylindrical and weakly spiraling, ending with a small, spoon-shaped thickening. Single medial ovary.

#### 18. *Microstomum inexcultus* sp. nov.; Fig. 5

Type: Asexual specimen; SMNH-Type-XXXX; GenBank Accession XXX Type Locality: Sweden, Saltö, 58°52’29’’N 11°08’41’’ E,10 cm depth, Habitat: Marine, eulittoral sand

GenBank Accession:

18S – XXX CO1 – XXX

Body length 0.85, 2 zooids. Colorless and without pigmented eyespots. Anterior end pointed; posterior tapered to a blunt tip with numerous adhesive papillae. Rhabdites absent. Ciliary pits clearly present. Preoral intestine long, stretching almost to anterior body end.

Reproductive system unknown.

#### 19. *Microstomum jenseni* Riedel, 1932

Synonyms:

*Microstomum tortipenis* Steinböck, 1938
Type: Deposition not recorded Type locality: Engelskmandshavn, Godhavn, Disko Island, Davis Strain Habitat: Marine benthic, to 300 m
Steinböck 1933 – Iceland, Isafjördur, fine sand, shallow water Westblad 1953, 1954 – Scandinavian west coast, Gullmar Fjord, 60-80 m depth, mud; Herdla, Kærnepollen, 40 m, mud Faubel 1974, 1977 – Germany, North Frisian Islands, Sylt Faubel and Warwick 2005 – United Kingdom, Scilly, St. Martin and surrounding

Body length around 0.6 mm. Colorless or whiteish with red-yellow intestine. Anterior conically pointed; posterior end long and tail-like with adhesive papillae. Rhabdites evenly distributed over body. Rhammite glands colored red at anterior end. Ciliary pits deep and narrow. Preoral intestine extending to level of brain. Intestine lightly ciliated or unciliated, with minotian cells. Nematocysts present.

Paired testes rounded to club-shaped. Paired vasa deferentia separately connected to vesicula granulorum. Male stylet 80 μm, shaped as an elongate weak spiral. Ovary unpaired

#### 20. *Microstomum laurae* Atherton & Jondelius 2018; Fig. 18B

Type: Asexual, SMNH-Type-8903, GenBank accession MF185712

Type location: Strömstad, Saltö, 58°52’29’’N, 11°08’41’’E, 10 cm depth

Habitat: Marine eulittoral

GenBank Accession

18S – XXX CO1 – MF185712-MF185713

Body length to 760 μm, 2 zooids. Body colorless and reflective of orange-red intestine. Anterior and posterior ends rounded. Posterior rim with approximately twenty adhesive papillae. Paired red eyespots 43 μm long, located in the lateral margins at level of the brain. Ciliary pits very small and shallow, directly below eyespots. Rhabdites concentrated in the anterior end above the pharynx. Nematocysts present. Preoral intestine extending anterior to brain.

Reproductive system unknown.

#### 21. *Microstomum lineare* (Müller, 1773); Fig. 17A

**Fig. 17.**
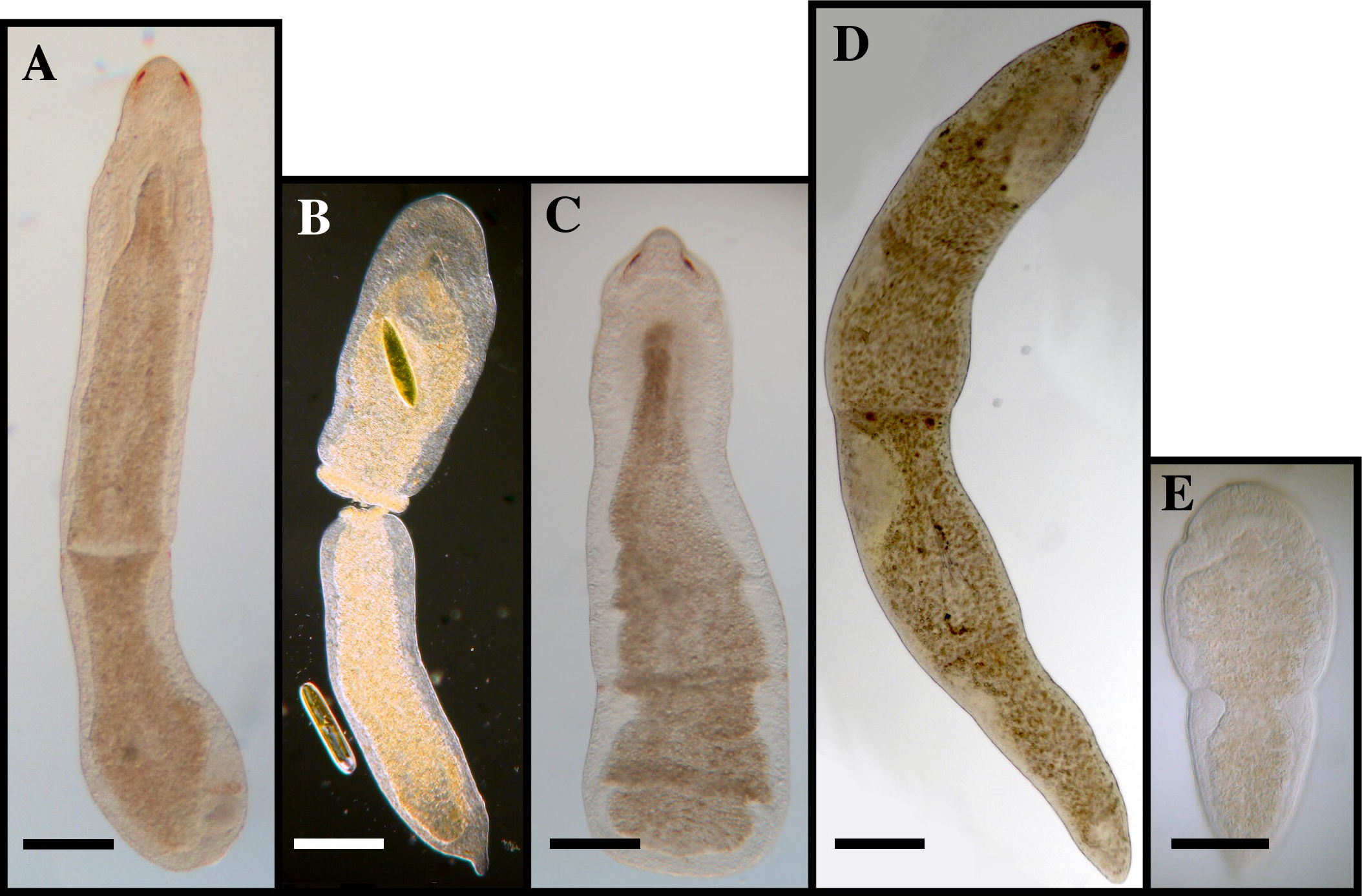
Examples of limnic *Microstomum*. A. *M. lineare* B. *M. zicklerorum* C. *M. tchaikovskyi* D. *M. artoisi* E. *M. bispiralis*. Bar = 200 μm

Synonyms:

*Fasciola linearis* – Müller, 1773, pg 67 *Planaria linearis* – Müller, 1776, pg 223 *Planaria vulgaris* – Fabricius, 1826, pg 18 *Derostoma flavicans* – Ehrenberg 1831 *Derostoma lineare* – Duges, 1828, pg 141 *Microstoma lineare* – Örsted 1843, pg 566 *Smigrostoma littoral* – Örsted 1845, pg 417 *Anotocelis linearis* – Diesing, 1862, pg 237 *Anotocelis flavicans* – Diesing, 1862, pg 242 *Microstomum caudatum* – Leidy 1852, pg 350 *Microstomum inerme* – Zacharias, 1894, pg 83 *Microstomum littorale* – Diesing 1861, pg 242

Type: No deposition recorded

Type Locality: Unknown

Habitat: Freshwater and brackish water to 7‰ salinity; littoral benthal or phytal; to 55+ m depth

An Der Lan 1939, 1961 – Macedonia, Lake Ohrid, springs, marshy banks, mud, to 33 m depth 1962 – Hungary, Danube

Atherton & Jondelius 2018b – Sweden, Finland, Belgium, benthic or phytal

Ax 1957 – Germany, Elbe, littoral

Bauchhenss 1971 – Germany, Franconia, Nuremberg and surrounding areas

Beklemishev 1921 – Russia, Saint Petersburg

Braccini & Leal-Zanchet 1913 – Brazil

Braun 1885 – Latvia, Estonia

Brusa 2006 – Argentina, Buenos Aires, Atalaya and Punta Piedras, benthic and phytal of rivers

Chodorowski 1959 – Poland, Harsz Lake

Dorner 1902 – Poland (Prussia)

Ferguson et al. 1939 – United States, Virginia, Mountain Lake Biological Station

Fuhrmann 1894 – Switzerland, Basel

Fulinski & Szynal 1932 – Ukraine, Poland, Lviv, Gródek

Gamo & Noreña 1998 – Spain, Iberian Peninsula

Gieysztor 1926 – Poland, Warsaw, Drozdowice, Zacisze

Graff 1911 – United States, Canada, Lake Ontario, on algae in puddles

1913 – United States, New Jersey, Michigan, Pennsylvania; Denmark

Hartog 1977 – Ireland, Cork, Bennet’s Lough

Heitkamp 1979, 1982 – Germany, Lower Saxony, ponds between Göttingen and Weser

Hofsten 1907 – Switzerland, Bern

1911 – Switzerland

Kaiser 1967, 1969 – Germany, Mühlhausen

1974 – Germany, Thuringia

Kolasa 1979, 1983 – Poland, Lake Zbe[chy, surrounding rivers and streams

Kolasa et al. 1987 – United States, New York, streams around Millbrook

Korgina, 2011 – Russia, Uvod’ Reservoir

Kosler 1962 – Germany, Hiddensee, benthic

Kraus 1965 – Austria, Neusiedlersee, Seewinkel National Park

Luther 1960 – Sweden, Denmark, Norway, lakes, rivers, Baltic sea, vegetation, sand, mud, to 30+ m

Mack-Fira◻ 1968 – Romania, Scurtu, Filipoiu Canal

1973, 1974 – Romania, Lagoon Complex Razelm-Sinoë

Meixner 1915 – Austria, Lunzer See

Müller & Faubel 1993 – Germany, Elbe Estuary

Nasonov 1919a, b, 1924 – Russia, Saint Petersburg and surrounding

1925 – Russia, Kola Peninsula

1929 – Japan

Noreña 1995 – Argentina, Paraná River

Noreña et al. 1999 – Spain, Extramadura

2006 – Peru, Loreto, Pacaya-Samiria National Reserve

2007 – Spain, Iberian Peninsula, Galician and Cantabrian coast

Okugawa 1953 – Japan

Papi 1952 – Italy, Pisa and surrounding areas, artificial ditches, still or slow waters

Petrovna & Vladimirovich, 2013 – Russia, Tartarstan, Kazan, Kaban Lake

Pörner 1966 – Germany, Jena and surrounding

Reisinger 1933 – Indonesia, Sumatra

Reisinger & Steinböck 1925 – Austria, Wörthersee

Rixen 1961 – Germany, Schleswig-Holstein, Lakes and ponds, phytal region, detritus-rich sand

1968 – Germany, Bodensee, benthic and phytal regions, to 50 m

Rogozin, 2016 – Russia, Miass, Il’menskoye

Ročník 2007 – Czechia, lednické rybníky ponds

Sabussow 1900 – Russia, Solowetzki Island

Schwank 1976 – Germany, Bondensee

Seifert 1939 – Germany, Hiddensee

Sibiriakova 1929 – Russia, Angara

Southern 1936 – Ireland, Cork, Tipperary, Wicklow

Steinböck 1933 – Faeroe Islands

1949, 1951a, b – Italy, Lago Maggiore, to 54 m

Tokinov & Berdnik, 2016 – Russia, Volga, Blue Lakes

Tu 1934 – China, Beijing, Tsing Hus University, campus pond

Urban-Malinga 2011 – Poland, Puck Bay, Hel Marine Station, 7 m depth

Valkanov 1926 - Bulgaria

Vara & Leal-Zanchet 2013 – Brazil, Rio Grande do Sul, in irrigated rice fields

Woodworth 1896 – United States, Michigan

Young 1970 – United Kingdom

1973 – United Kingdom, Wales

Zykov 1902 – Russia

GenBank Accession

28S – KC869844, specimen from United States; KP730557, KP730520, specimens from Finland; AJ270172 specimen from United Kingdom

18S – MH221198-257, MH221271-4, specimens from Sweden; KP730484, MH221258-70 specimens from Finland; MH221235-7, specimens from Belgium; D85092, specimen from Japan; KC869791, specimen from United States; U70082 collection location not listed 16S – KC869750, specimen from United States

ITS – MH221390-463, MH221475-6, specimens from Sweden; MH221464-474, specimens from Finland; MH221428-30, specimens from Belgium

CO1 – MH221368-80, KP730567, specimens from Finland; MF185697-9, MH221288-367, MH221381-2 specimens from Sweden; MH221329-31, specimens from Belgium; AY228756, AJ405979-80, specimens from United Kingdom Ef1a – AF288068, collection location not listed

Body length of solitary animal 0.4-1 mm; chains to 8 mm, 18 zooids maximum. Anterior end conically pointed. Posterior end tail-like. Body color variable: clear, grey, yellow, pink or brownish. Pigmented eyespots present as paired red stripes at dorso-anterior end. Amount of pigmentation highly variable and may be absent in some individuals based on light intensity.

Sensory bristles present. Cilia covering body, shorter in mouth. Ciliary pits at level of brain. Intestine ciliated and often including food remnants, including Cladocera, chironomid larvae, Copepoda, hydroids, Nematoda and other small animals. Nematocysts occuring throughout body.

Following TEM an SEM observations by Reuter (1978), groups of 6-8 adhesive papillae, each to 4 μm long, present in row at ventrolaterally at frontal end.

Nervous system includes brain, three pairs of nerve chords and pharyngeal ring system. Protonephridia with bilateral stems and pores in anterior body.

Male reproductive system with bilateral testes, circular or eliptical vesicular seminalis with prostate glands and separate gonopore. Male stylet spiraled, to 200 μm long. Female system with single ovary, ciliated oviduct and ventral pore.

Proterandrous development.

Notes: *Microstomum lineare* is an extremely common and abundant species in brackish and freshwater habitats worldwide and can be found February through late November or early December (Bauchhenss, 1971; Heitkamp, 1982). Lifecycle includes 4 generations per year, with sexual reproduction occurring primarily in late summer and fall. *M. lineare* purportedly has a global distribution and extremely wide tolerance for differing environmental conditions, including salinity (freshwater to 7‰; Karling 1974), temperature (to 47°C; Graff 1913), calcium (to 50 mg/L; Young 1973), oxygen and pH (Kaiser 1969; Heitkamp 1979, 1982).

Positive identification of *M. lineare* should be cautiously done since several other species of *Microstomum* are either very similar or identical in appearance to *M. lineare*. In part this is due to the very wide morphological variability reported for the species itself. Three species, *M. artoisi, M. tchaikovskyi*, and *M. zicklerorum*, were separated from *M. lineare* primarily through DNA evidence based on the mitochondrial cytochrome c oxidase subunit I and nuclear internal transcribed spacers I and II and 5.8S rDNA gene loci, and thus have few or no morphologically distinguishing characters (Atherton & Jondelius 2018b). A fourth, *M. giganteum*, though separable based on morphological evidence, can easily be misidentified and has historically been challenged as a “safe species”. However, *M. giganteum* is a larger species than *M. lineare* (8 zooid body length of ≥8.5 mm compared with 4-5 mm, respectively), without an elongated tail, adhesive papillae or pigmented eyespots, with a clearly visible esophagus, with pharyngeal glands restricted to the oral cavity only and with a three-part stylet (Kostenko 1988; Rogozin 2015).

#### 22. *Microstomum lotti* sp. nov.; Fig. 6

Type: MTP LS 660

Type Locality: Italy, Sant Andrea Bay, 42°48’31’’ N, 10°08’30’’ E

Habitat: Marine

GenBank Accession:

18S – KP730483, KP730493

Body length 0.8 mm, 2 zooid. Colorless and without pigmented eyespots. Anterior end conical; posterior end a spatulate tail plate. Adhesive papillae present in a row ventrally at the frontal region ~30 μm from anterior end, along posterior half of the body and very numerous at tail plate. Ciliary pits large and circular. Preoral intestine short. Rhabdites present.

Reproductive system unknown.

#### 23. *Microstomum lucidum* (Fuhrmann, 1896)

Synonyms:

*Microstoma lucidum* Fuhrmann 1896, pg 1011 *Microstomum hamatum* Westblad 1953 *Microstomum hanatum* WORMS (accessed 10 Aug. 2018)

Type: deposition not recorded Type location: English Channel, Baie de la Forêt Habitat: Marine sublittoral

Steinböck 1931 – Faroes, Vaagfjord, Suderø, mud 10 m Meixner 1938 – United Kingdom, Plymouth, Wembury Westblad 1953, 1954 – Norway, Bergen University, black mud; England, Plymouth, 40 m black mud; Blyth, 10 m, black mud. Karling 1966 – Norway, Herdla, Oöy and Blomöy

Body length to 1.5 mm, 4 zooids. Colorless and transparent with yellowish intestines. After Westblad (1953) may have (±) dark grey pigment spots in some individuals. Eyes absent. Ends bluntly rounded. Posterior end with adhesive papillae. Rhabdites present all over the body. Ciliary pits weakly visible. Preoral intestine extending to brain.

One or two testes, with only one being large and well developed, connecting via a single short vas deferens to the large vesicula seminalis. Stylet 60-70 μm, funnel shaped, proximally slightly thickened and curved like a hook distally. Antrum masculinum narrow.

Female system with unpaired ovary.

#### 24. *Microstomum marisrubri* sp. nov.; Fig. 7

Type: MTP LS 524

Type Locality: Egypt, Mangrove Bay, 25°52’15’’ N, 34°25’04’’ E Habitat: Marine

GenBank Accession:

18S – KP730505 CO1 – KP730580

Body length 0.65, 1 zooid. Body colorless, without pigmented eyespots. Body anterior to brain long, with rounded end. Posterior end blunt with numerous adhesive papillae. Adhesive papillae sparsely present over entire body. Ciliary pits present at level between anterior pharynx and brain. Preoral intestine short. Rhabdites absent. Nematocysts intensely concentrated at posterior end.

Male reproductive system including a single testis connected by short vas deferens to a rounded vesicula granulorum. Male stylet 63 μm long, shaped as a slightly tapering curved tube with a slight widening at the distal tip. Genital pores separate.

#### 25. *Microstomum melanophthalmum* Steinböck 1933

Type: deposition not recorded

Type location: Croatia, Rovinj, S. Andrea

Habitat: Marine, sublittoral to 7 m depth

Riedl, 1956, 1959 – Croatia, Isola Saint Andrea; Italy, Capo di Sorrento, Sicily, Porto Venere; France, Banyuls-sur-mer

Body length to 2.0 mm, to two zooids. Clear with yellow intestine. Black or dark brown eyes with distinct crescent lens near the frontal end. Anterior tip blunt. Posterior end conically rounded with adhesive tubes. Ciliary pits present just anterior to brain. Preoral intestine large, extending to well above brain. Rhabdites present.

Male reproductive system with seminal vesicular attached to stylet, 30 μm long, shaped as a sharply decreasing flat funnel connected to a tube hooked at an angle of 90° with a pointed end and terminal opening. Female system not seen.

#### 26. *Microstomum mortenseni* Riedel 1932

Type: deposition not recorded

Type location: Greenland, Disco Bay

Habitat: Marine, sublittoral benthal to 300 m depth

Body to 1.2 mm in solitary zooid. Based on Riedel’s (1932) original drawings, anterior end conically rounded and posterior end rounded and wide. Ciliary pits present at frontal end. Red color with yellow tinged rhammite glands. Pigmented eyespots absent. Rhabdites present at lateral parts of body. Preoral intestine long, anterior to brain. Intestine non-ciliated. Adhesive papillae not recorded.

Reproductive incompletely known. Ovary single with medial female gonopore. S shaped stylet.

#### 27. *Microstomum mundum* Graff 1905

Type: Deposition not recorded.

Type locality: Black Sea, Sevastopol Bay, St. Georges Monastery

Habitat: Marine, benthic in the sand

Body to 2 mm, 8 zooids. Colorless without pigmented eyespots. Anterior end conical. Adhesive papillae present in the posterior quarter of each well-developed zooid. Ciliary pits present at level of brain. Rhabdites present throughout the body, 16-20 μm long. Nematocysts present. Intestine, including preoral intestine, with characteristic lateral bulges and smaller lobules. Preoral intestine extending to a level halfway between the ciliary pits and anterior tip.

Reproductive system unknown.

##### -- *Microstomum ornatum* Uljanin 1870

Species dubiae by Westblad (1953) based on its incomplete description, perfunctory drawings, the posterior positioning of the ovary, which is unique among species of *Microstomum*, and the anterior direction of egg development, which seems to conflict with the posterior position of the female duct.

#### 28. *Microstomum papillosum* Graff 1882; Fig. 18D

**Fig. 18.**
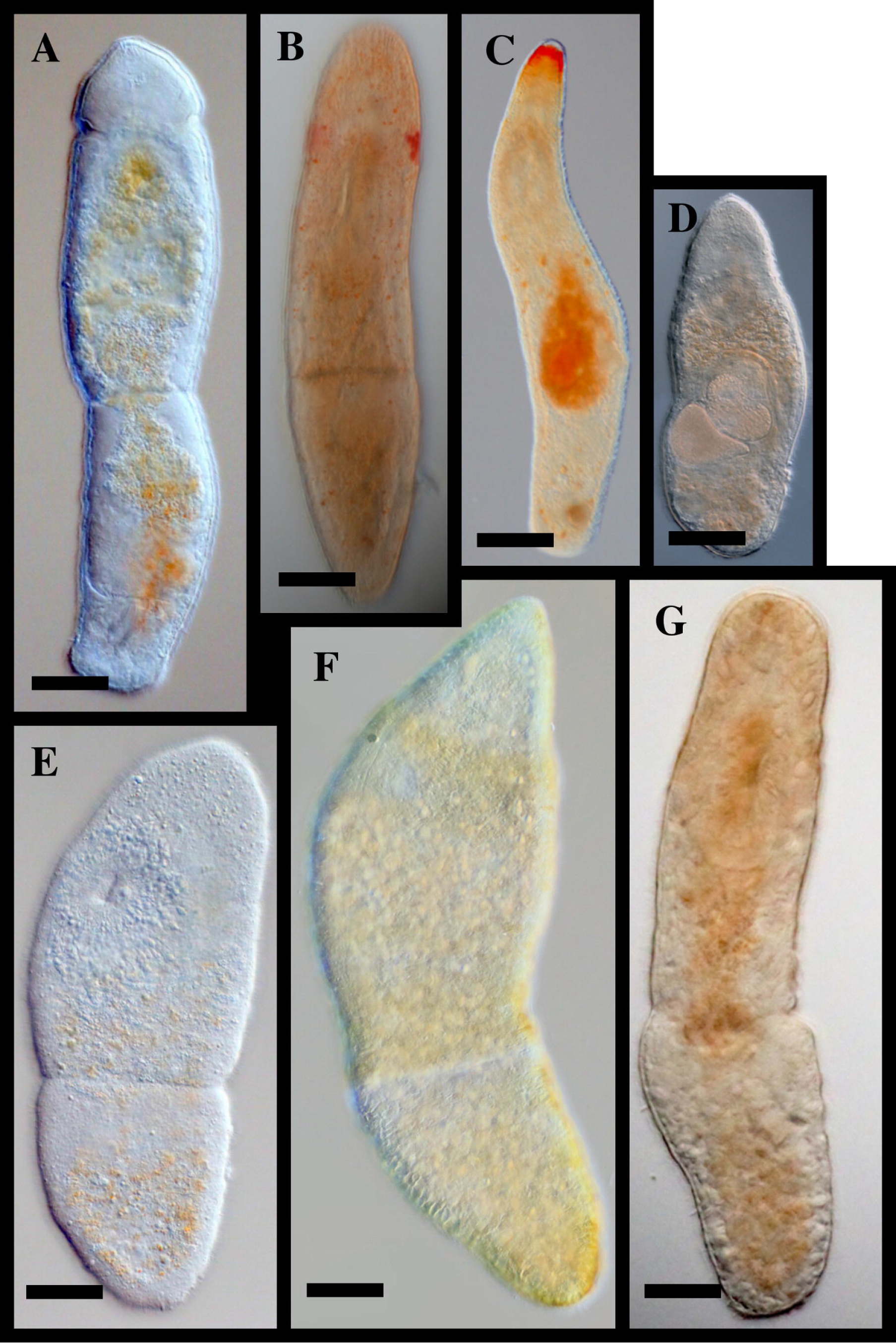
Examples of marine *Microstomum*. A. *M. edmondi* B. *M. laurae* C. *M. rubromaculatum* D. *M. papillosum* E. *M. crildensis* F. *M. compositum* G. *M. septentrionale.* Bar = 100 μm

Synonyms:

*Microstoma papillosum* – Claparède 1861, p152 Type: Deposition not recorded Type Locality: Sartor Oë, Norway Habitat: Marine, littoral to sublitteral. Armonies 1987 - North Sea, Sartorø near Bergen 2017 – Germany, Sylt, North Sea Ax 1959 – Black Sea, Bosporus, Marmara Sea Böhmig 1889 Faubel 1974, 1977 – Germany, Sylt, North Sea Faubel & Warwick 2005 – United Kingdom, Scilly, St. Martin and surrounding Hellwig 1987 Micoletzky 1910 – Gulf of Trieste, Croatia, Italy Riedl 1956 – Italy, Capo di Sorrento, Sicily, Porto Venere; France, Banyuls-sur-mer Steinböck 1938 – Iceland, Isafjördur, benthic and in algae Wehrenberg & Reise 1985 – Germany, Sylt, North Sea

GenBank Accession

28S – FJ715316, AF021330 specimens from Germany
18S – FJ715296, specimen from Germany
16S – KP730453, specimen from Germany
CO1 – KP730570, specimen from Germany

Body length to 1 mm, 5 zooids. Body shape with rounded anterior and a slightly tapered, rounded posterior. Body colorless except for reddish-brown intestine. Pigmented eyespots absent. Shallow ciliary pits present at level of anterior pharynx. Adhesive papillae scattered across the whole body but most numerous at anterior and posterior ends. Rhabdites present, especially at anterior end. Preoral intestine long, extending almost to anterior end.

Male genital system consisting of unpaired testis and vesicula seminalis. Stylet shaped as a curved tube, tapering to a thin distal terminal opening, 80 μm long. Female system including single ovary.

Notes: *Microstomum papillosum* has a somewhat similar stylet shape as *M. gabriellae* and *M. crildensis*, although its stylet is larger (80 μm compared to 50 and 32 μm, respectively) and tapers to a thinner tube distally. Other differences include, for *M. crildensis*, the size of the preoral intestine and the ciliary pits and the absence of rhabdites; and for *M. gabriellae*, the red body coloring and red eyespots.

#### 29. *Microstomum paradii* (Duges 1828)

Synonyms:

*Derostoma squalus* – Duges 1828, pg 142
Type: Deposition not recorded
Type locality: Romania, Transylvania, Cluj-Napoca
Habitat: Freshwater
Graff 1913 – France, Montpellier

Body to 2.5 mm, 2 zooids. Anterior end rounded; posterior tail-like with adhesive tubules. Colorless with light brown intestine. Black eyespots with conical lenses present anterior to ciliary pits. Rhabdites sparsely present throughout the body. Preoral intestine long.

Reproductive system not seen.

#### 30. *Microstomum proliferum* (Busch 1851)

Synonyms:

*Alaurina prolifera* Busch 1851, pg 114
Types: deposition not recorded
Type locality: United Kingdom, Skye Islands
Habitat: Marine, pelagic
Fewkes 1883 – USA, Newport
Graff 1913 – USA, New Jersey, Newport; Spain, Malaga

Body to 2.2 mm, 2 zooids. Pigmented eyes present lateral and posterior to mouth. Anterior end with conically pointed proboscis, non-ciliated, with many circular folds and denoted at the base by a tuft of cilia on each side. Sensory bristles present evenly spaced along lateral margins of the body, twice as long as the cilia, and single even longer bristle present at posterior end. Adhesive papillae present along entire body.

Notes: Brinkmann (1905) argued that this species is identical to *Alaurina composita* since any differences were either errors or derived from outdated methods of investigation. Graff (1913) argued against this, saying the two species differ in the sensory cilia and the location of the pigmented eyespots.

#### 31. *Microstomum punctatum* Dorner 1902

Types: Deposition not recorded
Type location: Linkenen Pond
Habitat: Freshwater pond, muddy with plants
Kaiser 1974 – Germany, Mühlhausen, Bad Langensalza
Wegelin 1966 – Germany, Halle, Elbe and surrounding

Body length 1 mm, 4 zooids maximum. Anterior end spade-like, pointed and separated from the posterior by a distinct constriction at level of the mouth. Posterior end tail-like. Body color brownish yellow, with numerous black dots at the anterior. Ciliary pits weak, at level of mouth. Adhesive papillae very sparsely present. Rhabdites absent. Pharynx very large, ¼ to ⅓ body size. Preoral intestine extremely small, without extending to anterior end of pharynx.

Single ovary. Male system unknown.

#### 32. *Microstomum rhabdotum* Marcus 1950

Type: Deposition not recorded
Type location: Brazil, São Sebastião
Habitat: Marine, between *Sargassum stenophyllum*

Body length to 0.7 mm, 3 zooids. Rounded anterior and posterior ends. Colorless and without pigmented eyespots. Adhesive papillae present at anterior, mouth, dividing zones, posterior. Biparte pharynx separated by annular fold with larger anterior cilia than posterior. Paired stripes of rhabdite bundles on the dorsolateral sides, joining at level of ciliary pits; median dorsal and ventral areas free of rhabdites; Preoral intestine reaches to anterior border of brain.

Reproductive system not seen.

#### 33. *Microstomum rubromaculatum* (Graff 1882); Fig. 18C

Synonyms:

*Microstoma rubromaculatum* Graff 1882, pg 251
Type: deposition not recorded
Type location: Gulf of Naples, Tyrrhenian Sea
Habitat: Marine, sublittoral phytal in *Sargassum* spp, benthic on shell sand.
Atherton & Jondelius 2018a – Sweden, Fiskebäckskil
Graff 1913 – France, Concarneau
Steinböck 1931 – Faroes, Vaagfjord, Suderø, mud 10 m
Westblad, 1934 – Norway, Herdla
Southern 1936 – Ireland, Dublin, Malahide Inlet, New Harbor, Galaway, 1-2 m
Steinböck 1938 – Iceland, Ísafjör*∂*ur, fine sand and algae, 1-2 m
Meixner 1938 – Enlgand, Wembury
Westblad 1953 – Sweden, Gullmar Fjord, Fiskebäckskil, 1-2 m

GenBank Accession:

18S – XXX CO1 – MF185684-MF185696, specimens from Sweden

Body length to 2 mm; maximum of 8 zooids. Very large red eyespot pigmentation, merging dorsally to form a broad transverse band at the anterior of the animal. Otherwise colorless or slightly pinkish reflective of intestines. Body ends bluntly rounded. Adhesive tubules present and concentrated at anterior and posterior ends. Rhabdites present throughout animal. Ciliary pits present with long cilia. Preoral intestine small.

Reproductive system based on specimens collected from Fiskebäckskil, Sweden (Atherton & Jondelius, 2018a). Male system with paired testes, copulatory apparatus, antrum masculinum and separate gonopore. Stylet 75-112 μm long, shaped as a single wide spiral bent around a 90° angle, terminating with a fingerlike hook. Female system with a single ovary situated mid-body, antrum, and a separate gonopore. Eggs develop caudally.

Notes: The amounts of eyespot pigmentation may be greatly variable, with pigmentation ranging from a bright circular band around the anterior end to two clearly distinct separate spots. Steinböck (1938) found one animal with a single large pigment spot in the middle that became thinner towards the margins of the body.

#### 34. *Microstomum schultei* sp. nov.; Fig. 8

Type: MTP LS 700
Type Locality: Italy, Fetovia Bay, 42°43’36’’ N, 10°09’33’’ E
Habitat: Marine

GenBank Accession:

18S – KP730494

Body length 1.5 mm, 1 zooid. Body light reddish-brown without pigmented eyespots. Anterior end rounded, with a small constriction at the ciliary pits. Posterior end conically blunted. Adhesive papillae and rhabdites present, concentrated at body ends. Ciliary pits present posterior to level of brain. Preoral intestine short.

Reproductive system with curved male stylet, tapering from a wider base to a narrow tube with a beveled tip.

#### 35. *Microstomum septentrionale* (Sabussow 1900); Fig. 18G

Synonyms:

*Microstoma ornatum* Sabussow 1897, pg 14 *Microstoma septentrionale* Sabussow 1900, pg 15, 81, 181
Type: Deposition not recorded
Type location: Russia, Kola Peninsula
Habitat: Marine, littoral among algae
Karling 1934 – Iceland, Herdla Kærnepollen
Westblad 1953 – Sweden, Gullmarfjord, Gåsövik
This paper – Sweden, Fiskebäckskil, 58°14′59′′ N, 11°26′45′′ E

GenBank Accession:

18S – XXX
CO1 – XXX

Maximum body length 1 mm, 2 zooids. Body translucent yellow with bright yellow intestines. Some specimens may have darker yellow or orange yellow in the anterior end. Pigmented eyespots absent. Anterior end rounded with a tapering rounded posterior end. Adhesive papillae sparsely scattered over the entire body. Preoral intestine reaching to level of anterior brain. Rhabdites present throughout the body, and especially concentrated in anterior end. Ciliary pits present at level of brain.

Single ovary and bilobed testis. Copulatory organ a curved tube, with a wide base and tapering distally to a point. Stylet length to 68 μm. Antrum masculinum present just anterior to stylet, large and ciliated. Separate ventral male and female pores.

#### 36. *Microstomum spiculifer* Faubel 1974

Type: Sagittal section series; Deposition numbers not listed
Type location: North Sea, North Frisian Uslands, Sylt Island, List
Habitat: Marine, eulittoral, semi-exposed sandy beach

Body length to 1.5 mm long, maximum 4 zooids. Body color clear with reddish brown intestine. Eyes absent. Ends rounded, posterior with multiple adhesive papillae. Preoral intestine reaching to level of brain or slightly anterior. Ciliated pits present at brain. Rhabdites present across body, concentrated at anterior.

Testis unpaired, located laterally and posterior to male copulatory apparatus. Masculine antrum elongated. Stylet strait, ca. to 55 μm long, decreasing from a width of 7-8 μm to a distal point. Ovary unpaired. Male and female pores separate.

#### 37. *Microstomum spiriferum* Westblad 1953

Lectotype: ◻, whole mount, SMNH-Type-XXXX; E. Westblad leg.**

Type Locality: Sweden, Gullmarsfjorden, Gullmarsvik

Habitat: Marine, mud, sublittoral between 10-60 m depth

Westblad 1953 – Sweden west coast, 10 m; Norway, Tromsö, Ramfjord, 10-25 m; United Kingdom, Scotland, Millport, 30-35 m

Body to 1.5 mm, generally composed of 2 zooids. Color pale reddish-yellow due to rich pigment cells in dorsal side. Red eyespots located at level of ciliary pits, may extend dorsally in some animals. Anterior and posterior ends rounded. Rhabdites present, concentrated in anterior and posterior ends. Adhesive papillae absent. Preoral intestine extending to level of ciliary pits.

Testis singular, small, connecting via a seminal duct to the vesicula seminalis and posterior penis stylet. Penis stylet large, 140 μm, with 5-6 dexiotropal torsions. With two female ducts and orifices, both connected to the ovary. The anterior acts as a ciliated oviduct and is separated from the ovary by a sphincter. The posterior acts as a female copulatory organ and may contain spermatozoids in the proximal end.

**A type specimen was not previously designated for *M. spiriferum.* However, fourteen original specimens were deposited at the Swedish Museum of Natural History, and thus, following the rules of the International Code of Zoological Nomenclature, we hereby designate one whole mount hermaphroditic specimen (formerly SMNH94968, now SMNH-Type-XXXX) as lectotype for *Microstomum spiriferum* WESTBLAD 1953. The remaining thirteen specimens (SMNH94966-7, SMNH94969-73, SMNH97839-44) are thus designated as paralectotypes.

#### 38. *Microstomum tchaikovskyi* Atherton & Jondelius 2018; Fig. 17C

Type: Asexual, SMNH-Type-9025; GenBank accession 18S MH221277, CO1 MH221385, ITS MH221479

Type Locality: Sweden, Sollentuna, 59°26’26’’N 18°00’03’’E, 15 cm depth Habitat: Freshwaters of Sweden and Finland, phytal to 20 cm depth

##### GenBank Accession

18S – MH221277, from Sweden; MH221278, from Finland

CO1 – MH221385, from Sweden; MH221386, from Finland

ITS – MH221479, from Sweden; MH221480, from Finland

##### Molecular Diagnosis

COI gene region with reference to GenBank Accession MH221288: 6-T; 15-A; T-16; 18-A; 30-G; 54-A; 69-A; 84-A; 180-G; 186-G; 228-A; 246-C; 279-A; 310-C; 312-T; 318-A; 321-A; 333-A; 351-A; 435-T; 486-G; 540-A; 603-A; 609-1; 625-C; 627-T

ITS1, 5.8S, ITS-2 gene region with reference to GenBank Accession MH221390: 121-T; 218-A; 288-T; 353-G; 358-G; 394-G; 497-T; 533-A; 730-G; 759-C; 768-C; 787-A; 800-T; 845-T; 887-T; 999-T

##### Morphological Description

Body length approximately 4 mm, 6 zooids. Body clear with darker intestines. Preoral intestine short. Anterior end abruptly tapering, posterior end tail-like without adhesive papillae. Sensory bristles present. Ciliary pits present at the level of anterior pharynx. Red eye pigmentation in two stripes, located dorsally at anterior end. May be weak in some individuals. Nematocysts few, scattered across the body.

Reproductive system not recorded.

Notes: See notes of *M. lineare*.

#### 39. *Microstomum trichotum* Marcus 1950

Lectotype: ♂, whole mount, SMNH-Type-XXXX, E. Marcus leg.**
Type Locality: Brazil, São Sebastião
Habitat: Marine, among calcareous algae incl. Corallinaceae and *Jania rubens*

Body length small, approximately 0.5 mm, 3 zooids. Body color white, without pigmented eyespots. Anterior end tapered and long, resembling the proboscis characteristic of *Alaurina* and densely ciliated, posterior end rounded with ca. 20 adhesive papillae. Preoral intestine to anterior brain. Ciliated pits small. Anterior rhammite tract absent. Rhabdites present throughout body.

Male reproductive system with single testis. Stylet shaped as a curved funnel, approximately 12 μm long. Female system not seen.

**A type specimen was not previously designated for *M. trichotum.* However, four original specimens were deposited at the Swedish Museum of Natural History, and thus, following the rules of the International Code of Zoological Nomenclature, we hereby designate one whole mount male specimen (formerly SMNH108151, now SMNH-Type-XXXX) as lectotype for *Microstomum trichotum* MARCUS 1950. The remaining three specimens (SMNH108149-50, SMNH10852) are thus designated as paralectotypes.

#### 40. *Microstomum ulum* Marcus 1950

Lectotype: Asexual, whole mount, SMNH-Type-XXXX, Leg. E. Marcus
Type locality: Brazil, São Sebastião
Habitat: Marine, sublittoral in coarse sand

Body size to 1.1 mm, 4 zooids. Colorless and without pigmented eyespots. Anterior end conically pointed, posterior separated by a constriction, forming a distinctly spatulate tail-plate. Adhesive papillae present. Rhabdites present throughout the body, but heavily concentrated at anterior end above the pharynx. Deep ciliary pits present at level of brain. Preoral intestine short.

Reproductive system not seen.

**A type specimen was not previously designated for *M. ulum.* However, three original specimens were deposited at the Swedish Museum of Natural History, and thus, following the rules of the International Code of Zoological Nomenclature, we hereby designate one whole mount asexual specimen (formerly SMNH108153, now SMNH-Type-XXXX) as lectotype for *Microstomum ulum* MARCUS 1950, while the remaining two specimens (SMNH108154-5) are thus designated as paralectotypes.

#### 41. *Microstomum weberi* sp. nov.; Fig. 9

Type: MTP LS 666
Type Locality: Italy, Pianosa, 42°34’29’’ N, 10°03’59’’ E
Habitat: Marine

GenBank Accession:

18S – KP730487 CO1 – KP730576

Body length of 0.65 mm, 2 zooid. Colorless and without pigmented eyespots. Anterior end conical; posterior end widely blunt. Adhesive papillae present at posterior end and in a ventral row at the frontal end 25 μm from anterior tip. Ciliary pits bottle-shaped. Preoral intestine extending to anterior brain or just beyond. Large rhabdites bundles present at anterior and posterior ends.

Reproductive system unknown.

#### 42. *Microstomum westbladi* nom. nov.; Fig. 10

Synonym:

*Microstomum papillosum* GRAFF 1882, sensu Westblad 1953, pg 406-7
Type: Asexual, sectioned, SMNH-Type-XXXX Type Locality: Norway, Herdla Habitat: Marine, sublittoral sand and mud

GenBank Accession:

18S – XXX CO1 – XXX

Westblad, 1953 – Norway, Herdla, Kværnepoll current; Sweden, Gullmarfjord This paper – Sweden, Malmön, 58°20’18’’ N 11°20’26’’ E

Body length to 2.0 mm, 5 zooids. Colorless and without pigmented eyespots. Anterior and posterior ends bluntly rounded; posterior with numerous adhesive papillae. Ciliary pits small. Rhabdites numerous and intensely concentrated at frontal end. Adhesive papillae present sparsely along body entire length, with characteristic grouping present at anterior tip: field longest at both lateral sides and curving upwards toward the middle. Preoral intestine extending to brain.

Reproductive system including male stylet, 60 μm long, and shaped as a smooth curve 180° curve, with a maximum width ~⅙ of the way to the distal end, decreasing proximally and also distally to a point.

#### 43. *Microstomum zicklerorum* Atherton & Jondelius 2018; Fig. 17B

Type: Asexual, SMNH-Type-9026; GenBank accession 18S MH221281, CO1 MH221389, ITS MH221483

Type Locality: USA, Massachusetts, North Chelmsford 42°38’13’’N 71°23’30’’W Habitat: Freshwater benthic or phytal; to 25 cm depth

##### GenBank Accession

18S – MH221279 – MH221280
CO1 – MH221387 – MH221389
ITS – MH221481 – MH221483

##### Molecular Diagnosis

COI gene region with reference to GenBank Accession MH221288: 24-A; 30-T; 123-T; 165-A; 186-A; 192-G; 201-G; 267-C; 300-C; 318-G; 324-G; 330-A; 336-C; 387-C; 501-A; 525-G; 555-G; 558-A; 588-A; 609-G; 630-C

ITS1, 5.8S, ITS-2 gene region with reference to GenBank Accession MH221390: 197-A; 317-A; 507-T; 829-A; 879-A; 888-C; 1000-A

##### Morphological Description

Body length to 1.5 mm, 2 zooids. Body clear and reflective of intestines. Red eye pigmentation extremely weak or not present. Anterior end conical, posterior end tail-like without adhesive papillae. Sensory bristles present. Ciliary pits present with circular openings lateral at the level of mid-pharynx. Preoral intestine short. Mouth and pharynx large leading to strait intestine. Intestine containing desmid algae. Nematocysts present throughout the body. Reproductive system not recorded.

Notes: See notes of *M. lineare.*

### Gen. Myozonella Beklemishev 1955

Microstomidae with ciliary pits at frontal end and intestine extending anterior to mouth opening (preoral intestine). Muscle ring present surrounding intestine. Asexual reproduction through fissioning present; sexual reproduction unknown. Freshwater, one species.

Type species: *Myozonella microstomoides* BEKLEMISHEV 1955

#### 1. *Myozonella microstomoides* Beklemishev 1955

Type: Deposition not recorded

Type locality: Russia, Near Bryansk in town of Sudimir

Habitat: Freshwater, phytal in *Utriculara* sp and *Calopteryx* sp.

Body size to 1.0 mm, 2 zooids. Colorless and without pigmented eyespots. Anterior end conically pointed, posterior with thin tail. Adhesive papillae and rhabdites absent. Ciliary pits present dorsolaterally, narrow and deep. Preoral intestine extending to level of ciliary pits. Muscle ring present surrounding intestines just posterior to pharynx.

Reproductive system not seen.

## Acknowledgements

We would like to thank many people for their valuable help, insight and support: Dr. Tom Artois, Edmond ATHERTON, Laura Atherton, Adam Bates, Dr. Marco Curini-Galletti, Dr. Rick Hochberg, Ylva Jondelius and the entire Zickler family. We are grateful to the staffs of the Stensoffa Research Station, Abisko Scientific Research station, and Umeå Marina Forskningcentrum. We are also particularly indebted to Dr. Lukas Schärer for the collection, documentation and sequencing of four of the species included in this manuscript and for generally providing insights and instruction on Macrostomorpha taxonomy. Many thanks also to Cassandra Vion-Bailly for creating and providing all drawings in this manuscript. This research was supported by a grant from the Swedish Taxonomy Initiative (no. DHA 2014-151 4.3) to UJ.

---

**Fig. S1: Concatenated 18S and CO1 gene tree with all specimens**

**Table S1. Collecting information for the specimens used in this study**. Collecting location, specific coordinates and dates are given for each specimen, where available. In addition, the GenBank Accession Numbers for 18S, and CO1 sequences and references are listed.

**Table S2: Primer sequences, references and protocols for a mplification of 18S and CO1 sequences.**

**Table S3: A summary of the morphological characters for each species**.

